# Complete genome characterisation of a novel coronavirus associated with severe human respiratory disease in Wuhan, China

**DOI:** 10.1101/2020.01.24.919183

**Authors:** Fan Wu, Su Zhao, Bin Yu, Yan-Mei Chen, Wen Wang, Yi Hu, Zhi-Gang Song, Zhao-Wu Tao, Jun-Hua Tian, Yuan-Yuan Pei, Ming-Li Yuan, Yu-Ling Zhang, Fa-Hui Dai, Yi Liu, Qi-Min Wang, Jiao-Jiao Zheng, Lin Xu, Edward C. Holmes, Yong-Zhen Zhang

## Abstract

Emerging and re-emerging infectious diseases, such as SARS, MERS, Zika and highly pathogenic influenza present a major threat to public health^1–3^. Despite intense research effort, how, when and where novel diseases appear are still the source of considerable uncertainly. A severe respiratory disease was recently reported in the city of Wuhan, Hubei province, China. At the time of writing, at least 62 suspected cases have been reported since the first patient was hospitalized on December 12^nd^ 2019. Epidemiological investigation by the local Center for Disease Control and Prevention (CDC) suggested that the outbreak was associated with a sea food market in Wuhan. We studied seven patients who were workers at the market, and collected bronchoalveolar lavage fluid (BALF) from one patient who exhibited a severe respiratory syndrome including fever, dizziness and cough, and who was admitted to Wuhan Central Hospital on December 26^th^ 2019. Next generation metagenomic RNA sequencing^4^ identified a novel RNA virus from the family *Coronaviridae* designed WH-Human-1 coronavirus (WHCV).

Phylogenetic analysis of the complete viral genome (29,903 nucleotides) revealed that WHCV was most closely related (89.1% nucleotide similarity similarity) to a group of Severe Acute Respiratory Syndrome (SARS)-like coronaviruses (genus *Betacoronavirus*, subgenus *Sarbecovirus*) previously sampled from bats in China and that have a history of genomic recombination. This outbreak highlights the ongoing capacity of viral spill-over from animals to cause severe disease in humans.

---

Seven patients, comprising five men and two women, were hospitalized at the Central Hospital of Wuhan from December 14 through December 28, 2019. The median age of the patients was 43, ranging from 31 to 70 years old. The clinical characteristics of the patients are shown in Table 1. Fever and cough were the most common symptoms. All patients had fever with body temperatures ranging from 37.2°C to 40°C. Patients 1, 2, 5, 6 and 7 had cough, while patients 1, 2 and 7 presented with severe cough with phlegm at onset of illness. Patients 4 and 5 also complained of chest tightness and dyspnea. Patients 1, 3, 4 and 6 experienced dizziness and patient 3 felt weakness. No neurological symptoms were observed in any of the patients. Bacterial culture revealed the presence of *Streptococcus* bacteria in throat swabs from patients 3, 4 and 7. Combination antibiotic, antiviral and glucocorticoid therapy were administered. Unfortunately, patient 1 and 4 showed respiratory failure: patient 1 was given high flow noninvasive ventilation, while patient 4 was provided with nasal/face mask ventilation (Table 1).

**Table 1.** Clinical symptoms and patient data

| Characteristic | Patient 1 | Patient 2 | Patient 3 | Patient 4 | Patient 5 | Patient 6 | Patient 7 |
| --- | --- | --- | --- | --- | --- | --- | --- |
| Age (Year) | 41 | 44 | 42 | 70 | 31 | 51 | 43 |
| Sex | M | F | M | F | M | M | M |
| Date of illness onset | Dec 20,2019 | Dec 22,2019 | Dec 24,2019 | Dec 24,2019 | Dec 21,2019 | Dec 16,2109 | Dec 14,2019 |
| Date of admission | Dec 26,2019 | Dec 22,2019 | Dec 28,2019 | Dec 28,2019 | Dec 28,2019 | Dec 27,2019 | Dec 14,2019 |
| Fever | + | + | + | + | + | + | + |
| Body Temperature (°C) | 38.4 | 37.3 | 39 | 37.9 | 38.7 | 37.2 | 38 |
| Cough | + | + | + | + | + | + | + |
| Sputum Production | + | + | - | - | - | - | + |
| Dizzy | + | - | + | + | - | + | - |
| Weakness | + | - | + | - | - | - | - |
| Chest tightness | + | - | - | + | + | - | - |
| Dyspnea | + | - | - | + | + | + | - |
| Bacterial culture | - | - | streptococcus pneumoniae | streptococcus pneumoniae | - | - | streptococcus pneumoniae |
| Glucocorticoid therapy | No | No | Yes | Yes | Yes | Yes | No |
| Antibiotic therapy | Cefoselis | Ceftazidime, Levofloxacin | Cefminox | Cefminox, moxifloxacin | Cefminox | No | No |
| Antiviral therapy | Oseltamivir | No | Oseltamivir, ganciclovir | Oseltamivir, ganciclovir | Oseltamivir, ganciclovir | Oseltamivir, ganciclovir | No |
| Oxygen therapy | mechanical ventilation | No | No | Mask | No | No | No |

Epidemiological investigation by the Wuhan CDC revealed that all the suspected cases were linked to individuals working in a local indoor seafood market. Notably, in addition to fish and shell fish, a variety of live wild animals including hedgehogs, badgers, snakes, and birds (turtledoves) were available for sale in the market before the outbreak began, as well as animal carcasses and animal meat. No bats were available for sale. While the patients might have had contact with wild animals in the market, none recalled exposure to live poultry.

Patient 1 was a 41-year-old man with no history of hepatitis, tuberculosis or diabetes. He was admitted and hospitalized in Wuhan Central Hospital 6 days after the onset of illness. The patient reported fever, chest tightness, unproductive cough, pain and weakness for one week on presentation. Physical examination of cardiovascular, abdominal and neurologic examination was normal. Mild lymphopenia (less than 900 cells per cubic milli-meter) was observed, but white blood cell and blood platelet count was normal in a complete blood count (CBC) test. Elevated levels of C-reactive protein (CRP, 41.4 mg/L of blood, reference range 0-6 mg/L) was observed and levels of aspartate aminotransferase, lactic dehydrogenase, and creatine kinase were slightly elevated in blood chemistry tests. The patient had mild hypoxemia with oxygen levels of 67mmHg by the Arterial Blood Gas (ABG) Test. On the first day of admission (day 6 after the onset of illness), chest radiographs were abnormal with air-space shadowing such a ground-glass opacities, focal consolidation and patchy consolidation in both lungs (Figure 1). Chest computed tomographic (CT) scans revealed bilateral focal consolidation, lobar consolidation and patchy consolidation, especially in the lower lung. A chest radiograph revealed a bilateral diffuse patchy and fuzzy shadow on day 5 after admission (day 11 after the onset of illness). Preliminary aetiological investigation excluded the presence of influenza virus, *Chlamydia pneumoniae* and *Mycoplasma pneumoniae* by commercial pathogen antigen detection kits and confirmed by PCR. Other common respiratory pathogens, including adenovirus, were also negative by qPCR (Figure S1). The condition of the patient did not improve after three days of treatment with combined antiviral and antibiotic therapy. He was admitted to the intensive care unit (ICU) and treatment with a high flow non-invasive ventilator was initiated. The patient was transferred to another hospital in Wuhan for further treatment 6 days after admission.

**Figure 1.**
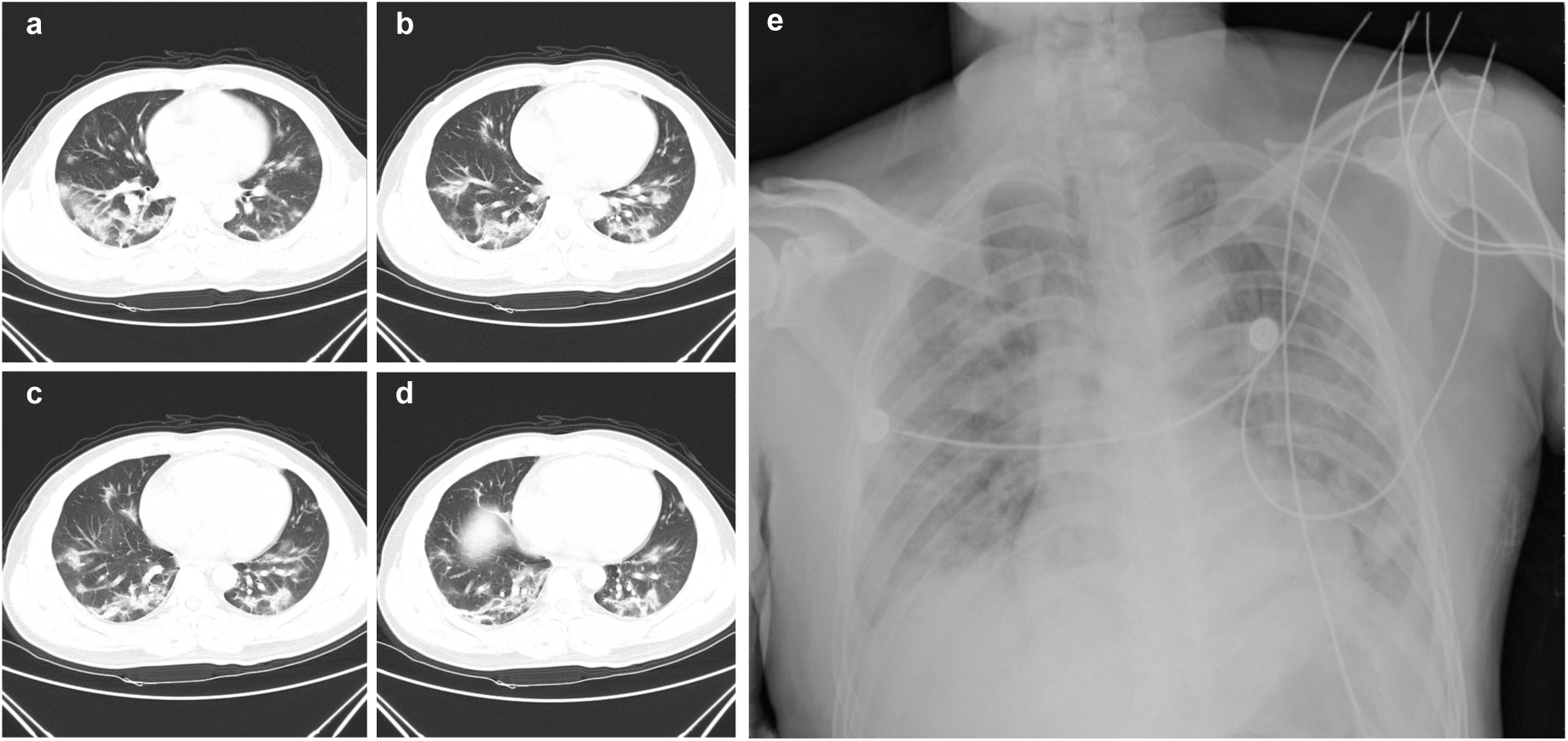
Chest radiographs of patient 1. **a. b. c. d.** Chest computed tomographic scans of Patient 1 were obtained on the day of admission (day 6 after the onset of illness). Bilateral focal consolidation, lobar consolidation, and patchy consolidation were clearly observed, especially in the lower lung. **e**. Chest radiograph of patient 1 was obtained on day 5 after admission (day 11 after the onset of illness). Bilateral diffuse patchy and fuzzy shadow were observed.

To investigate the possible aetiologic agents associated this disease, we collected bronchoalveolar lavage fluid (BALF) from patient 1 and performed deep meta-transcriptomic sequencing. All the clinical specimens were handled in a biosafety level 3 laboratory at the Shanghai Public Health Clinical Center. Total RNA was extracted from 200μl BAL fluid and a meta-transcriptomic library was constructed for pair-end (150 bp) sequencing using an Illumina MiniSeq as previously described^4–7^. In total, we generated 56,565,928 sequence reads that were *de novo* assembled and screened for potential aetiologic agents. Of the 384,096 contigs assembled by Megahit^8^, the longest (30,474 nucleotides [nt]) had high abundance and was closely related to a bat SARS-like coronavirus isolate - bat-SL-CoVZC45 (GenBank Accession MG772933) - previously sampled in China, with a nt identity of 89.1% (Table S1 and S2). The genome sequence of this novel virus, as well as its termini, were determined and confirmed by RT-PCR^9^ and 5’/3’ RACE kits (TaKaRa), respectively. This new virus was designated as WH-Human 1 coronavirus (WHCV) (and has also been referred to as ‘2019-nCoV’) and its whole genome sequence (29,903 nt) has been assigned GenBank accession number MN908947. Remapping the RNA-seq data against the complete genome of WHCV resulted in an assembly of 123,613 reads, providing 99.99% genome coverage at a mean depth of 6.04X (range: 0.01X-78.84X) (Figure S2). The viral load in the BALF sample was estimated by quantitative PCR (qPCR) to be 3.95×10^8^ copies/mL (Figure S3).

The viral genome organization of WHCV was characterized by sequence alignment against two representative members of the genus *Betacoronavirus*: a human-origin coronavirus (SARS-CoV Tor2, AY274119) and a bat-origin coronavirus (Bat-SL-CoVZC45, MG772933) (Figure 2). The un-translational regions (UTR) and open reading frame (ORF) of WHCV were mapped based on this sequence alignment and ORF prediction. The WHCV viral genome was similar to these two coronaviruses (Figure 2 and Table S3), with a gene order 5’-replicase ORF1ab-S-envelope(E)-membrane(M)-N-3’. WHCV has 5’ and 3’ terminal sequences typical of the betacoronaviruses, with 265 nt at the 5’ terminal and 229 nt at the 3’ terminal region. The predicted replicase ORF1ab gene of WHCV is 21,291 nt in length and contained 16 predicted non-structural proteins (Table S4), followed by (at least) 13 downstream ORFs. Additionally, WHCV shares a highly conserved domain (LLRKNGNKG: amino acids 122-130) with SARS-CoV in nsp1. The predicted S, ORF3a, E, M and N genes of WHCV are 3,822, 828, 228, 669 and 1,260 nt in length, respectively. In addition to these ORFs regions that are shared by all members of the subgenus *Sarbecovirus*, WHCV is similar to SARS-CoV in that it carries a predicted ORF8 gene (366 nt in length) located between the M and N ORF genes. The functions of WHCV ORFs were predicted based on those of known coronaviruses and given in Table S5. In a manner similar to SARS CoV Tor2, a leader transcription regulatory sequence (TRS) and nine putative body TRSs could be readily identified upstream of the 5’ end of ORF, with the putative conserved TRS core sequence appeared in two forms – the ACGAAC or CUAAAC (Table S6).

**Figure 2.**
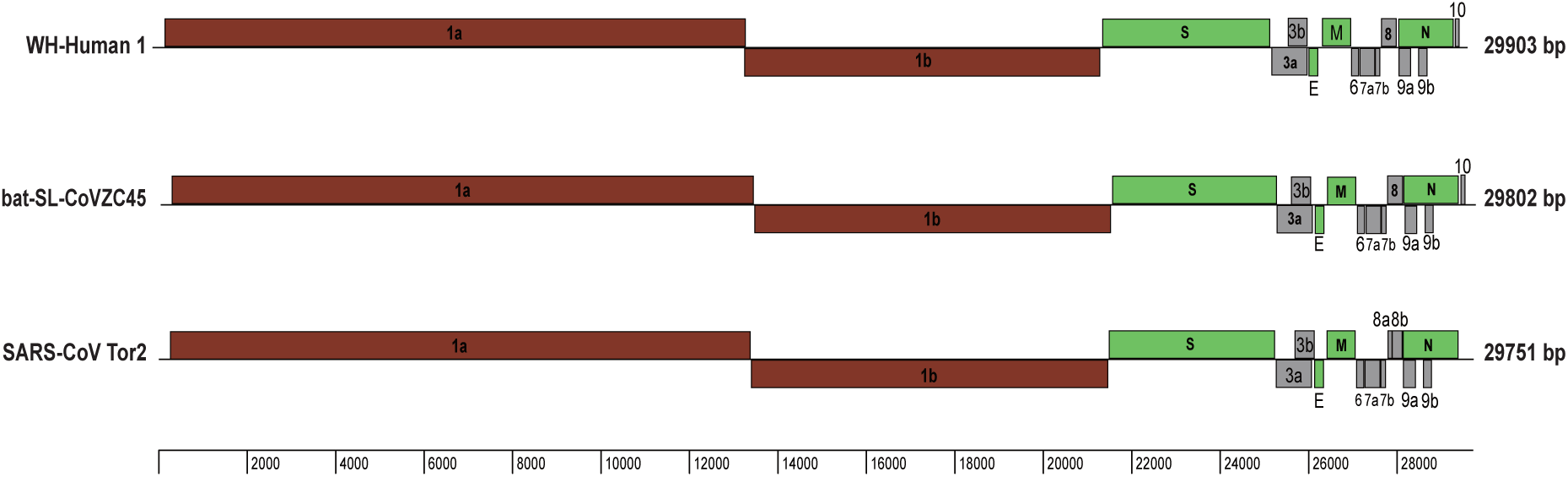
Genome organization of SARS and SARS-like CoVs including Tor2, CoVZC45 and WHCV determined here.

To determine the evolutionary relationships between WHCV and previously identified coronaviruses, we estimated phylogenetic trees based on the nucleotide sequences of the whole genome sequence, non-structural protein genes ORF1a and 1b, and the main structural proteins encoded by the S, E, M and N genes (Figures 3 and S4). In all phylogenies WHCV clustered with members of the subgenus *Sarbecovirus*, including the SARS-CoV responsible for the global SARS pandemic of 2002-2003^1, 2^, as well as a number of SARS-like coronaviruses sampled from bats. However, WHCV changed topological position within the subgenus *Sarbecovirus* depending on which gene was used, suggestive of a past history of recombination in this group of viruses (Figures 3 and S4). Specifically, in the S gene tree (Figure S4), WHCV was most closely related to the bat coronavirus bat-SL-CoVZC45 with 82.3% amino acid (aa) identity (and ∼77.2% aa identity to SARS CoV; Table S3), while in the ORF1b phylogeny WHCV fell in a basal position within the subgenus *Sarbecovirus* (Figure 3). This topological division was also observed in the phylogenetic trees estimated for conserved domains in the replicase polyprotein pp1ab (Figure S5).

**Figure 3.**
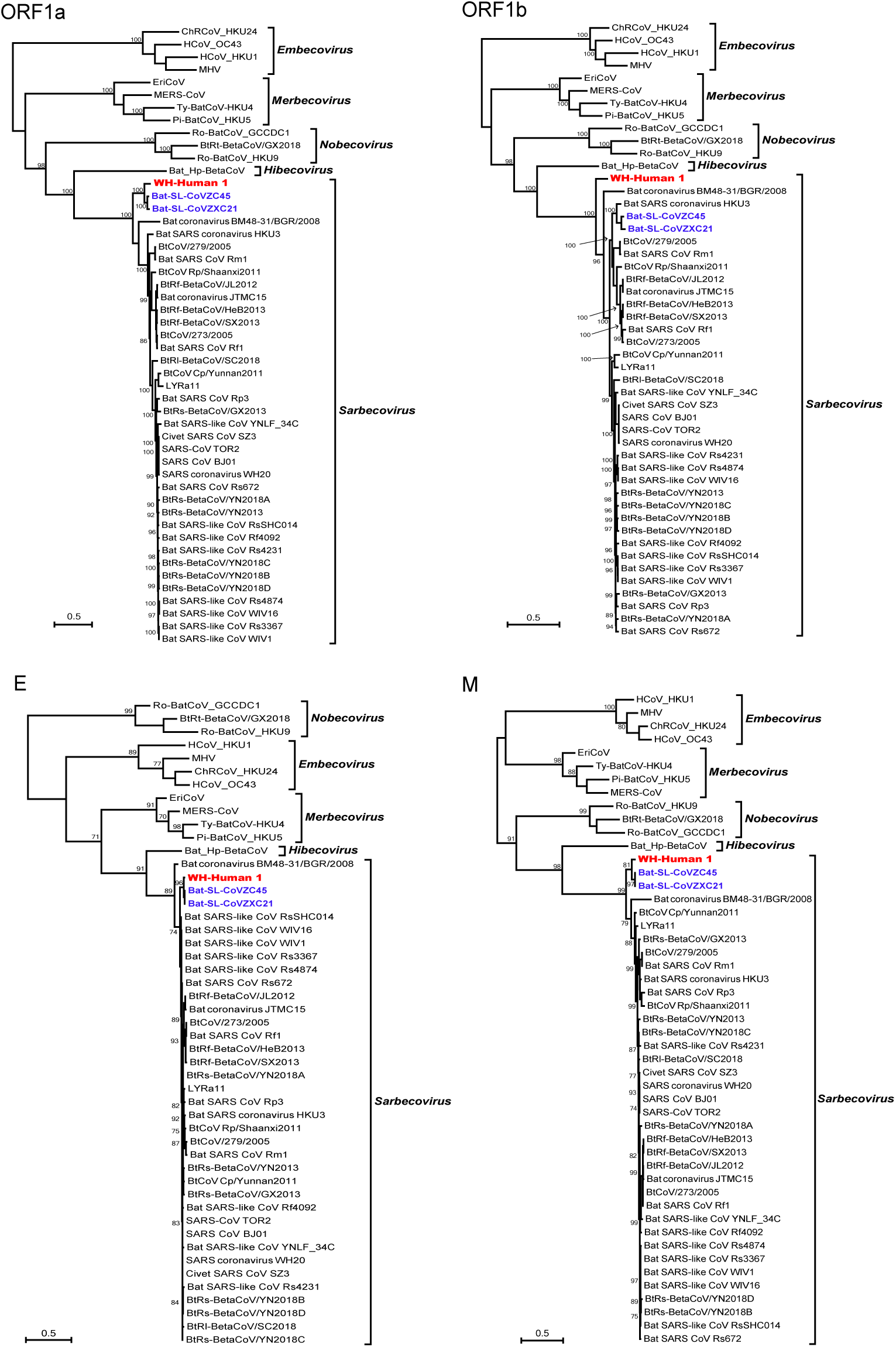
Maximum likelihood phylogenetic trees of nucleotide sequences of the ORF1a, ORF1b, E and M genes of WHCV and related coronaviruses. Numbers (>70) above or below branches indicate percentage bootstrap values for the associated nodes. The trees were mid-point rooted for clarity only. The scale bar represents the number of substitutions per site.

To better understand the potential of WHCV to infect humans, the receptor-binding domain (RBD) of its spike protein was compared to those in SARS-CoVs and bat SARS-like CoVs. The RBD sequences of WHCV were more closely related to those of SARS-CoVs (73.8%-74.9% aa identity) and SARS-like CoVs including strains Rs4874, Rs7327 and Rs4231 (75.9%-76.9% aa identity) that are able to use the human ACE2 receptor for cell entry (Table S7)^10^. In addition, the WHCV RBD was only one amino acid longer than the SARS-CoV RBD (Figure 4a). In contrast, other bat SARS-like CoVs including the Rp3 strain that cannot use human ACE2^11^, had amino acid deletions at positions 473-477 and 460-472 compared to the SARS-CoVs (Figure 4a). The previously determined^12^ crystal structure of SARS-CoV RBD complexed with human ACE2 (PDB 2AJF) revealed that regions 473-477 and 460-472 directly interact with human ACE2 and hence may be important in determining species specificity (Figure 4b). We predicted the three-dimension protein structures of WHCV, Rs4874 and Rp3 RBD domains by protein homology modelling using the SWISS-MODEL server and compared them to the crystal structure of SARS-CoV RBD domains (PDB 2GHV) (Figure 4, c-f). In accord with the sequence alignment, the predicted protein structures of WHCV and Rs4874 RBD domains were closely related to that of SARS-CoVs and different from the predicted structure of the RBD domain from Rp3. In addition, the N-terminus of WHCV S protein is more similar to that of SARS-CoV rather than other human coronaviruses (HKU1 and OC43) (Figure S6) that can bind to sialic acid^13^. In sum, the high similarities of amino acid sequences and predicted protein structure between WHCV and SARS-CoV RBD domains suggest that WHCV may efficiently use human ACE2 as a cellular entry receptor, perhaps facilitating human-to-human transmission^10, 14–15^.

**Figure 4.**
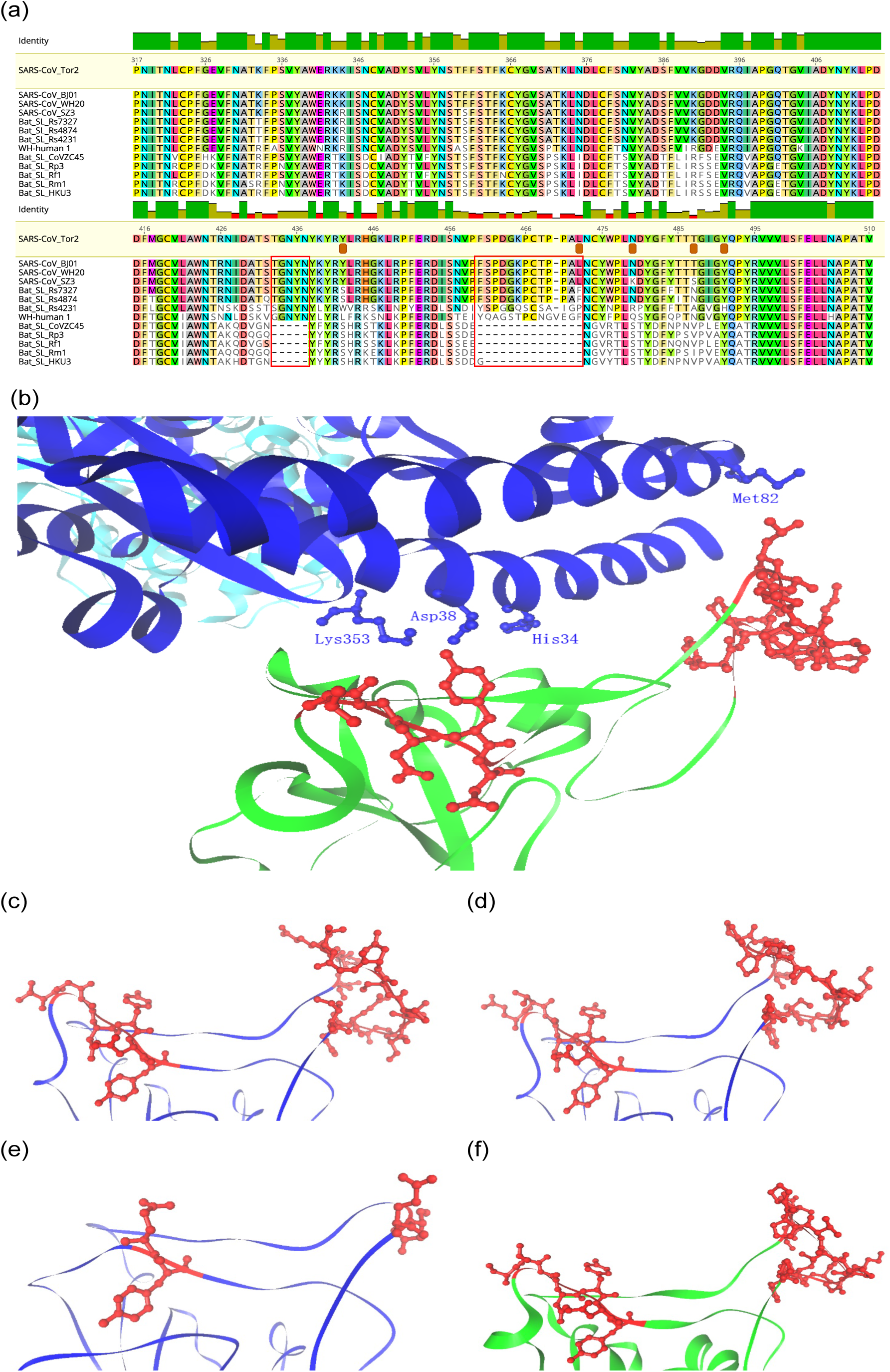
Analysis of receptor-binding domain (RBD) of the spike (S) protein of WHCV coronavirus. **(a)** Amino acid sequence alignment of SARS-like CoV RBD sequences. Three bat SARS-like CoVs, which could efficiently utilize the human ACE2 as receptor, had an RBD sequence of similar size to SARS-CoV, and WHCV contains a single Val 470 insertion. The key amino acid residues involved in the interaction with human ACE2 are marked with a brown box. In contrast, five bat SARS-like CoVs had amino acid deletions at two motifs (amino acids 473-477 and 460-472) compared with those of SARS-CoV, and Rp3 has been reported not to use ACE2.^11^ **(b)** The two motifs (aa 473-477 and aa 460-472) are shown in red on the crystal structure of the SARS-CoV spike RBD complexed with receptor human ACE2 (PDB 2AJF). Human ACE2 is shown in blue and the SARS-CoV spike RBD is shown in green. Important residues in human ACE2 that interact with SARS-CoV spike RBD are marked. **(c)** Predicted protein structures of RBD of WHCV spike protein based on target-template alignment using ProMod3 on the SWISS-MODEL server. The most reliable models were selected based on GMQE and QMEAN Scores. Template: 2ghw.1.A, GMQE: 0.83; QMEAN:-2.67. Motifs resembling amino acids 473-477 and 460-472 of the SARS-CoV spike protein are shown in red. **(d)** Predicted structure of RBD of SARS-like CoV Rs4874. Template: 2ghw.1.A, GMQE:0.99; QMEAN:-0.72. Motifs resembling amino acids 473-477 and 460-472 of the SARS-CoV spike protein are shown in red. (**e**) Predicted structure of the RBD of SARS-like CoV Rp3. Template: 2ghw.1.A, GMQE:0.81, QMEAN:-1.50. **(f)** Crystal structure of RBD of SARS-CoV spike protein (green) (PDB 2GHV). Motifs of amino acids 473-477 and 460-472 are shown in red.

To further characterize putative recombination events in the evolutionary history of the sarbecoviruses the whole genome sequence of WHCV and four representative coronaviruses - Bat SARS-like CoV Rp3, CoVZC45, CoVZXC21 and SARS-CoV Tor2 - were analysed using the Recombination Detection Program v4 (RDP4)^16^. Although the similarity plots suggested possible recombination events between WHCV and SARS CoVs or SARS-like CoVs (Figure S7), there was no significant evidence for recombination across the genome as a whole. However, some evidence for past recombination was detected in the S gene of WHCV and SARS CoV and bat SARS-like CoVs (WIV1 and RsSHC014) (p<3.147×10^-3^ to p<9.198×10^-9^), with similarity plots suggesting the presence of recombination break points at nucleotides 1,029 and 1,652 that separated the WHCV S gene into three regions (Figure 5). In phylogenies of the fragment nt 1 to 1029 and nt 1652 to the end of the sequence, WHCV was most closely related to Bat-SL-CoVZC45 and Bat-SL-CoVZXC21, whereas in the region nt 1030 to 1651 (the RBD region) WHCV grouped with SARS CoV and bat SARS-like CoVs (WIV1 and RsSHC014) that are capable of direct human transmission^14, 17^.

**Figure 5.**
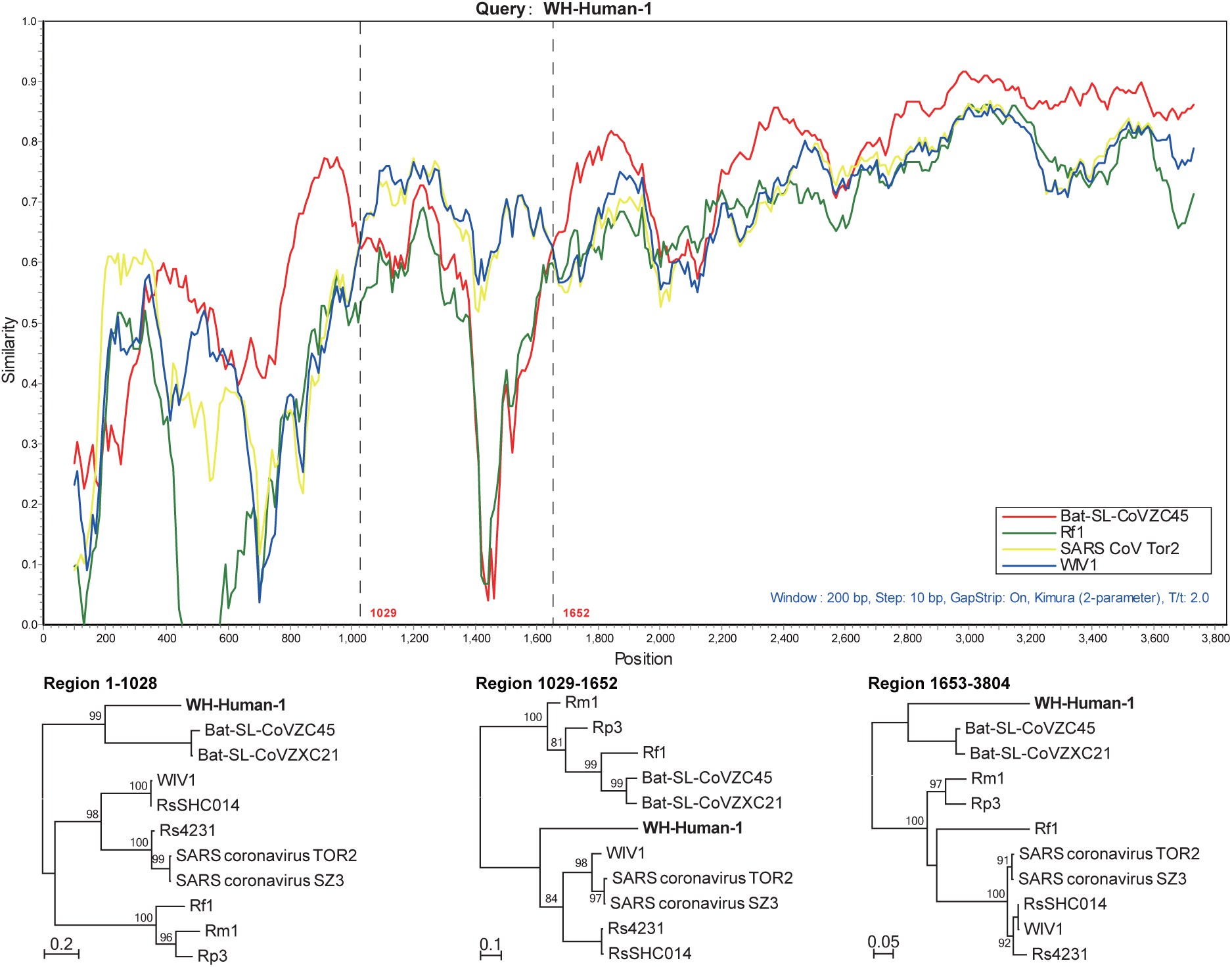
Possible recombination events in the S gene of sarbecoviruses. A sequence similarity plot (upper panel) reveals two putative recombination break-points shown by black dashed lines, with their locations indicated at the bottom. The plot shows S gene similarity comparisons of the WHCV (query) against SARS-CoV Tor2 and bat SARS-like CoVs WIV1, Rf1 and CoVZC45. Phylogenies of major parental region (1-1028 and 1653-3804) and minor parental region (1029-1652) are shown below the similarity plot. Phylogenies were estimated using a ML method and mid-point rooted for clarity only. Numbers above or below branches indicate percentage bootstrap values.

Coronaviruses are associated with a number of infectious disease outbreaks in humans, including SARS in 2002/3 and MERS in 2012^1, 18^. Four other coronaviruses - human coronaviruses HKU1, OC43, NL63 and 229E - are also associated with respiratory disease^19–22^. Although SARS-like coronaviruses have been widely identified in mammals including bats since 2005 in China^9, 23–25^, the exact origin of human-infected coronaviruses remains unclear. Herein, we describe a novel coronavirus - WHCV - in BALF from a patient experiencing severe respiratory disease in Wuhan, China. Phylogenetic analysis suggested that WHCV represents a novel virus within genus *Betacoronavirus* (subgenus *Sarbecovirus*) and hence that exhibits some genomic and phylogenetic similarity to SARS-CoV^1^, particularly in the RBD. These genomic and clinical similarities to SARS, as well as its high abundance in clinical samples, provides evidence for an association between WHCV and the ongoing outbreak of respiratory disease in Wuhan.

The identification of multiple SARS-like-CoVs in bats led to the idea that these animals act as the natural reservoir hosts of these viruses^19, 20^. Although SARS-like viruses have been identified widely in bats in China, viruses identical to SARS-CoV have not yet been documented. Notably, WHCV is most closely related to bat coronaviruses, even exhibiting 100% aa similarity to Bat-SL-CoVZC45 in the nsp7 and E proteins. Hence, these data suggest that bats are a possible reservoir host of WHCV. However, as a variety of animal species were for sale in the market when the disease was first reported, more work is needed to determine the natural reservoir and any intermediate hosts of WHCV.

## Acknowledgements

This study was supported by the Special National Project on investigation of basic resources of China (Grant SQ2019FY010009) and the National Natural Science Foundation of China (Grants 81861138003 and 31930001). ECH is supported by an ARC Australian Laureate Fellowship (FL170100022).

## Author Contributions

Y.-Z.Z. conceived and designed the study. S.Z, Y.H, Z.-W.T. and M.-L.Y. performed the clinical work and sample collection. B.Y and J.-H.T. performed epidemiological investigation and sample collection. F.W, Z.-G.S., L.X., Y.-Y.P., Y.-L.Z., F.-H.D., Y.L., J.-J.Z. and Q.-M.W. performed the experiments. Y.-M.C., W.W., F.W., E.C.H. and Y.-Z.Z. analysed the data. Y.-Z.Z. E.C.H. and F.W. wrote the paper with input from all authors. Y.-Z.Z. led the study.

## Competing interests

The authors declare no competing interests.

## Methods

### Cases and collection of clinical data and samples

Patients presenting with acute onset of fever (>37.5°C), cough, and chest tightness, and who were admitted to Wuhan Central Hospital in Wuhan city, China, were considered as suspected cases. During admission, bronchoalveolar lavage fluid (BALF) was collected and stored at −80°C until further processing. Demographic, clinical and laboratory data were retrieved from the clinical records of the confirmed patients.

### RNA library construction and sequencing

Total RNA was extracted from the BALF sample of patient 1 using the RNeasy Plus Universal Mini Kit (Qiagen) following the manufacturer’s instructions. The quantity and quality of the RNA solution was assessed using a Qbit machine and an Agilent 2100 Bioanalyzer (Agilent Technologies) before library construction and sequencing. An RNA library was then constructed using the SMARTer Stranded Total RNA-Seq Kit v2 (TaKaRa, Dalian, China). Ribosomal RNA (rRNA) depletion was performed during library construction following the manufacturer’s instructions. Paired-end (150 bp) sequencing of the RNA library was performed on the MiniSeq platform (Illumina). Library preparation and sequencing were carried out at the Shanghai Public Health Clinical Center, Fudan University, Shanghai, China.

### Data processing and viral agent identification

Sequencing reads were first adaptor- and quality-trimmed using the Trimmomatic program^26^. The remaining reads (56,565,928 reads) were assembled *de novo* using both the Megahit (version 1.1.3)^8^ and Trinity program (version 2.5.1)^27^ with default parameter settings. Megahit generated a total of 384,096 assembled contigs (size range: 200-30,474 nt), while Trinity generated 1,329,960 contigs with a size range of 201 to 11,760 nt. All of these assembled contigs were compared (using blastn and Diamond blastx) against the entire non-redundant nucleotide (Nt) and protein (Nr) database, with *e-*values set to 1×10^-10^ and 1×10^-5^, respectively. To identify possible aetiologic agents present in the sequence data, the abundance of the assembled contigs was first evaluated as the expected counts using the RSEM program^28^ implemented in Trinity. Non-human reads (23,712,657 reads), generated by filtering host reads using the human genome (human release 32, GRCh38.p13, downloaded from Gencode) by Bowtie2^29^, were used for the RSEM abundance assessment.

As the longest contigs generated by Megahit (30,474 nt) and Trinity (11,760 nt) both had high similarity to the bat SARS-like coronavirus isolate bat-SL-CoVZC45 and were at high abundance (Table S1 and S2), the longer one (30,474 nt) that covered almost the whole virus genome was used for primer design for PCR confirmation and genome termini determination. Primers used in PCR, qPCR and RACE experiments are listed in Table S8. The PCR assay was conducted as described previously^9^ and the complete genome termini was determined using the Takara SMARTer RACE 5’/3’ kit (TaKaRa) following the manufacturer’s instructions. Subsequently, the genome coverage and sequencing depth were determined by remapping all of the adaptor- and quality-trimmed reads to the whole genome of WHCV using Bowtie2^29^ and Samtools^30^.

The viral loads of WHCV in BALF of patient 1 were determined by quantitative real-time RT-PCR with Takara One Step PrimeScript™ RT-PCR Kit (Takara RR064A) following the manufacturer’s instructions. Real-time RT-PCR was performed using 2.5μl RNA with 8pmol of each primer and 4pmol probe under the following conditions: reverse transcription at 42°C for 10 minutes, and 95°C for 1 minute, followed by 40 cycles of 95°C for 15 seconds and 60°C for 1 minute. The reactions were performed and detected by ABI 7500 Real-Time PCR Systems. PCR product covering the Taqman primers and probe region was cloned into pLB vector using the Lethal Based Simple Fast Cloning Kit (TIAGEN) as standards for quantitative viral load test.

### Virus genome characterization and phylogenetic analysis

For the newly identified virus genome, the potential open reading frames (ORFs) were predicted and annotated using the conserved signatures of the cleavage sites recognized by coronavirus proteinases, and were processed in the Lasergene software package (version 7.1, DNAstar). The viral genes were aligned using the L-INS-i algorithm implemented in MAFFT (version 7.407)^31^.

Phylogenetic analyses were then performed using the nucleotide sequences of various CoV gene data sets: (i) Whole genome, (ii) ORF1a, (iii) ORF1b, (iv) nsp5 (3CLpro), (v) RdRp (nsp12), (vi) nsp13 (Hel), (vii) nsp14 (ExoN), (viii) nsp15 (NendoU), (ix) nsp16 (O-MT), (x) spike (S), and the (xi) nucleocapsid (N). Phylogenetic trees were inferred using the Maximum likelihood (ML) method implemented in the PhyML program (version 3.0)^32^, using the Generalised Time Reversible substitution (GTR) model and Subtree Pruning and Regrafting (SPR) branch-swapping. Bootstrap support values were calculated from 1,000 pseudo-replicate trees. The best-fit model of nucleotide substitution was determined using MEGA (version 5)^33^. Amino acid identities among sequences were calculated using the MegAlign program implemented in the Lasergene software package (version 7.1, DNAstar).

### Genome recombination analysis

Potential recombination events in the history of the sarbecoviruses were assessed using both the Recombination Detection Program v4 (RDP4)^16^ and Simplot (version 3.5.1)^34^. The RDP4 analysis was conducted based on the complete genome (nucleotide) sequence, employing the RDP, GENECONV, BootScan, maximum chi square, Chimera, SISCAN, and 3SEQ methods. Putative recombination events were identified with a Bonferroni corrected p-value cut-off of 0.01. Similarity plots were inferred using Simplot to further characterize potential recombination events, including the location of breakpoints.

### Analysis of RBD domain of WHCV spike protein

An amino acid sequence alignment of WHCV, SARS-CoVs, bat SARS-like CoVs RBD sequences was performed using MUSCLE^35^. The predicted protein structures of the spike protein RBD were estimated based on target-template alignment using ProMod3 on SWISS-MODEL server (https://swissmodel.expasy.org/). The sequences of the spike RBD domains of WHCV, Rs4874 and Rp3 were searched by BLAST against the primary amino acid sequence contained in the SWISS-MODEL template library (SMTL, last update: 2020-01-09, last included PDB release: 2020-01-03). Models were built based on the target-template alignment using ProMod3. The global and per-residue model quality were assessed using the QMEAN scoring function^36^. The PDB files of the predicted protein structures were displayed and compared with the crystal structures of SARS-CoV spike RBD (PDB 2GHV)^37^ and the crystal of structure of SARS-CoV spike RBD complexed with human ACE2 (PDB 2AJF)^12^.

## Supplementary legends

### Supplementary Tables

**Table S1.** The top 50 abundant assembled contigs generated using the Megahit program.

**Table S2.** The top 80 abundant assembled contigs generated using the Trinity program.

**Table S3.** Amino acid identities of the selected predicted gene products between the novel coronavirus (WHCV) and known betacoronaviruses.

**Table S4.** Cleavage products of the replicase polyproteins of WHCV.

**Table S5.** Predicted gene functions of WHCV ORFs.

**Table S6.** Coding of potential and putative transcription regulatory sequences of the genome sequence of WHCV.

**Table S7.** Amino acid identities of the RBD sequence between SARS- and bat SARS-like CoVs.

**Table S8.** PCR primers used in this study.

### Supplementary Figures

**Figure S1.** Detection of other respiratory pathogens by qPCR.

**Figure S2.** Mapped read count plot showing the coverage depth per base of the WHCV genome.

**Figure S3.** Detection of WHCV in clinical samples by RT-qPCR. **(a)** Specificity of the WHCV primers used in RT-qPCR. Test samples comprised clinical samples that are positive for at least one of the following viruses: Influenza A virus (09H1N1 and H3N2), Influenza B virus, Human adenovirus, Respiratory syncytial virus, Rhinovirus, Parainfluenza virus type 1-4, Human bocavirus, Human metapneumovirus, Coronavirus OC43, Coronavirus NL63, Coronavirus 229E and Coronavirus HKU1. **(b-c)** Standard curve. **(d)** Amplification curve of WHCV.

**Figure S4.** Maximum likelihood phylogenetic trees of the nucleotide sequences of the whole genome, S and N genes of WHCV and related coronaviruses. Numbers (>70) above or below branches indicate percentage bootstrap values. The trees were mid-point rooted for clarity only. The scale bar represents the number of substitutions per site.

**Figure S5.** Maximum likelihood phylogenetic trees of the nucleotide sequences of the 3CL, RdRp, Hel, ExoN, NendoU, and O-MT genes of WHCV and related coronaviruses. Numbers (>70) above or below branches indicate percentage bootstrap values. The trees were mid-point rooted for clarity only. The scale bar represents the number of substitutions per site.

**Figure S6.** Amino acid sequence comparison of the N-terminal domain (NTD) of spike protein of WHCV, bovine coronavirus (BCoV), mouse hepatitis virus (MHV) and human coronavirus (HCoV OC43 and HKU1) that can bind to sialic acid and the SARS-CoVs that cannot. The key residues^13^ for sialic acid binding on BCoV, MHV, HCoV OC43 and HKU1 were marked with a brown box.

**Figure S7.** A sequence similarity plot of WHCV, SARS- and bat SARS-like CoVs revealing putative recombination events.

