## Supplementary Tables and Figures for "Complete genome characterisation of a novel coronavirus associated with severe human respiratory disease in Wuhan, China"

#### Supplementary Table

**Table S1.** The top 50 abundant assembled contigs generated using the Megahit program.

| Contigs | Length | Abundance [a] | Result of blast against Nt database | | | Result of blast against Nr database | | |
|---|---|---|---|---|---|---|---|---|
| | | | Blast hit | Identity (%) | e-value | Blastx hit | Identity (%) | e-value |
| k141_7767 | 1549 | 1355719.87 | CP019721 Veillonella parvula strain UTDB1-3, complete genome | 99.613 | 0 | CUP36263.1 Uncharacterised protein [Bacteroides xylanisolvens] | 62.6 | 2.30E-32 |
| k141_261129 | 274 | 1170229.79 | CP022041 Prevotella melaninogenica strain FDAARGOS_306 chromosome 2, complete sequence | 100 | 9.55E-140 | CUO90010.1 Uncharacterised protein [Prevotella copri] | 96.9 | 6.90E-09 |
| k141_357441 | 391 | 633828.63 | CP016205 Prevotella scopos JCM 17725 strain W2052 chromosome 2 genome | 94.359 | 8.16E-167 | KDS36881.1 hypothetical protein M091_0855 [Parabacteroides distasonis str. 3776 D15 i] | 53.2 | 1.30E-21 |
| k141_132235 | 2540 | 598061.73 | CP019721 Veillonella parvula strain UTDB1-3, complete genome | 97.338 | 0 | ABP91180.1 unknown protein [Streptococcus suis 98HAH33] | 63.8 | 6.60E-53 |
| k141_10046 | 240 | 497112.72 | CP022041 Prevotella melaninogenica strain FDAARGOS_306 chromosome 2, complete sequence | 100 | 6.52E-121 | CUO89876.1 Cell wall-associated hydrolase [Prevotella copri] | 96.2 | 2.00E-36 |
| k141_246896 | 239 | 461925.98 | CP022040 Prevotella melaninogenica strain FDAARGOS_306 chromosome 1, complete sequence | 99.582 | 1.09E-118 | EHG15578.1 hypothetical protein HMPREF9138_01799, partial [Prevotella histicola F0411] | 95.7 | 8.70E-16 |
| k141_245870 | 569 | 429581.29 | CP023863 Prevotella jejuni strain CD3:33 chromosome I, complete sequence | 97.88 | 0 | EHG15578.1 hypothetical protein HMPREF9138_01799, partial [Prevotella histicola F0411] | 93.6 | 2.70E-15 |
| k141_165136 | 556 | 390619.28 | CP023863 Prevotella jejuni strain CD3:33 chromosome I, complete sequence | 99.64 | 0 | KWW26465.1 hypothetical protein AUK64_2547 [bacterium P201] | 62.5 | 4.10E-32 |
| k141_170961 | 241 | 389549.4 | HM322133 Uncultured bacterium clone ncd392h10c1 16S ribosomal RNA gene, partial sequence | 100 | 1.82E-121 | EDM51784.1 hypothetical protein EUBVEN_00788 [Eubacterium ventriosum ATCC 27560] | 60 | 5.20E-08 |
| k141_72317 | 502 | 383496.8 | JQ459396 Uncultured bacterium clone 070027_126 16S ribosomal RNA gene, partial sequence | 99.452 | 0 | AOE06246.1 hypothetical protein [uncultured bacterium] | 56.8 | 2.00E-14 |
| k141_61109 | 353 | 304459.13 | AP018050 Prevotella melaninogenica DNA, complete genome, strain: GAI 07411 | 99.717 | 0 | EDM19151.1 hypothetical protein BACCAC_03785 [Bacteroides caccae ATCC 43185] | 71.3 | 2.60E-24 |
| k141_46290 | 765 | 257373.81 | HQ616399 Prevotella sp. ICM55 16S ribosomal RNA gene, partial sequence | 99.213 | 0 | EDO51672.1 hypothetical protein BACUNI_04219 [Bacteroides uniformis ATCC 8492] | 72.4 | 1.60E-23 |
| k141_39967 | 526 | 252641 | FJ557960 Uncultured bacterium clone ET_G_3d09 16S ribosomal RNA gene, partial sequence | 98.289 | 0 | AOE11686.1 hypothetical protein [uncultured bacterium] | 69.8 | 8.30E-19 |
| k141_56198 | 4633 | 225245.16 | CP012072 Actinomyces meyeri strain W712, complete genome | 95.006 | 0 | GAN11851.1 hydrolase, partial [Mucor ambiguus] | 51.3 | 7.00E-61 |
| k141_211996 | 245 | 224901.41 | CP022041 Prevotella melaninogenica strain FDAARGOS_306 chromosome 2, complete sequence | 98.776 | 1.12E-118 | EFI73306.1 cell wall-associated hydrolase [Prevotella bryantii B14] | 93.8 | 4.30E-26 |
| k141_248606 | 411 | 219591 | CP023864 Prevotella jejuni strain CD3:33 chromosome II, complete sequence | 100 | 0 | EFN91701.1 hypothetical protein HMPREF9018_1166 [Prevotella amnii CRIS 21A-A] | 95.5 | 3.70E-38 |
| k141_250867 | 334 | 206021.11 | LC356098 Uncultured bacterium 221MH06016 gene for 16S rRNA, partial sequence | 94.895 | 1.18E-144 | OUD19662.1 hypothetical protein BUN10_26130 [Vibrio parahaemolyticus] | 80.6 | 4.80E-20 |
| k141_228454 | 1444 | 199997.19 | JX424618 Prevotella sp. Sc00026 clone contig00026c genomic sequence | 89.646 | 0 | KWW26465.1 hypothetical protein AUK64_2547 [bacterium P201] | 76 | 7.60E-30 |
| k141_325767 | 474 | 198930.08 | CP022041 Prevotella melaninogenica strain FDAARGOS_306 chromosome 2, complete sequence | 99.789 | 0 | KWW24027.1 hypothetical protein F082_2040 [bacterium F082] | 60 | 2.90E-18 |
| k141_132017 | 646 | 190579.71 | LT906445 Veillonella parvula strain NCTC11810 genome assembly, chromosome: 1 | 100 | 0 | CKL43271.1 Cell wall-associated hydrolase [Neisseria meningitidis] | 74 | 6.80E-63 |
| k141_218984 | 499 | 188987.21 | AP018049 Prevotella melaninogenica DNA, complete genome, strain: GAI 07411 | 97.595 | 0 | EDY97039.1 hypothetical protein BACPLE_00421 [Bacteroides plebeius DSM 17135] | 86.2 | 9.30E-36 |
| k141_124227 | 287 | 186562.92 | AP018050 Prevotella melaninogenica DNA, complete genome, strain: GAI 07411 | 98.27 | 3.62E-139 | KWW26465.1 hypothetical protein AUK64_2547 [bacterium P201] | 63.2 | 4.00E-23 |
| k141_154714 | 309 | 183385.02 | CP023864 Prevotella jejuni strain CD3:33 chromosome II, complete sequence | 99.029 | 3.84E-154 | EFN91701.1 hypothetical protein HMPREF9018_1166 [Prevotella amnii CRIS 21A-A] | 94.2 | 8.00E-46 |

| | | | | | | | | |
|---|---|---|---|---|---|---|---|---|
| k141_225856 | 557 | 181751.03 | EF510660 Uncultured bacterium clone P2D11-613 16S ribosomal RNA gene | 100 | 0 | EDM19152.1 hypothetical protein BACCAC_03786 [Bacteroides caccae ATCC 43185] | 79.5 | 2.90E-54 |
| k141_281403 | 479 | 180948.65 | CP022041 Prevotella melaninogenica strain FDAARGOS_306 chromosome 2, complete sequence | 94.395 | 6.29E-144 | CUO90010.1 Uncharacterised protein [Prevotella copri] | 93.8 | 1.00E-07 |
| k141_246050 | 2139 | 179525.89 | CP003667 Prevotella sp. oral taxon 299 str. F0039 plasmid, complete sequence | 96.282 | 0 | EFC67102.1 LOW QUALITY PROTEIN: hypothetical protein HMPREF0670_02906, partial [Prevotella sp. oral taxon 317 str. F0108] | 77.8 | 7.50E-34 |
| k141_356852 | 558 | 175931.25 | FJ557960 Uncultured bacterium clone ET_G_3d09 16S ribosomal RNA gene, partial sequence | 100 | 8.01E-89 | EHG15578.1 hypothetical protein HMPREF9138_01799, partial [Prevotella histicola F0411] | 95.7 | 5.40E-16 |
| k141_2268 | 595 | 154024.63 | JQ459396 Uncultured bacterium clone 070027_126 16S ribosomal RNA gene, partial sequence | 99.138 | 1.63E-175 | AOE06246.1 hypothetical protein [uncultured bacterium] | 55.4 | 2.00E-13 |
| k141_9212 | 499 | 152834.88 | CP023863 Prevotella jejuni strain CD3:33 chromosome I, complete sequence | 94.567 | 0 | KWW24027.1 hypothetical protein F082_2040 [bacterium F082] | 68.4 | 8.20E-24 |
| k141_228442 | 359 | 151855.46 | LC359097 Uncultured bacterium 81AD08008 gene for 16S rRNA, partial sequence | 95.822 | 7.48E-162 | CDN41090.1 hypothetical protein BN871_AB_00880 [Paenibacillus sp. P22] | 67.8 | 1.20E-32 |
| k141_30268 | 326 | 146328.06 | CP019721 Veillonella parvula strain UTDB1-3, complete genome | 100 | 1.43E-168 | ETJ17454.1 hypothetical protein Q620_VSAC00705G0001, partial [Veillonella sp. DORA_A_3_16_22] | 97.9 | 3.30E-42 |
| k141_235167 | 283 | 144371.6 | LC359516 Uncultured bacterium 83MG01013 gene for 16S rRNA, partial sequence | 100 | 9.83E-145 | AOE11686.1 hypothetical protein [uncultured bacterium] | 77.3 | 1.40E-12 |
| k141_78882 | 1400 | 137390.81 | CP022386 Capnocytophaga gingivalis strain H1496 chromosome, complete genome | 99.5 | 0 | KWW27340.1 hypothetical protein AUK64_2223 [bacterium P201] | 78.6 | 7.80E-48 |
| k141_76959 | 3696 | 130539.21 | CP001685 Leptotrichia buccalis DSM 1135, complete genome | 93.051 | 0 | EEX74124.1 hypothetical protein GCWU000323_01827 [Leptotrichia hofstadii F0254] | 92.7 | 2.20E-49 |
| k141_123601 | 273 | 123375.65 | CP022041 Prevotella melaninogenica strain FDAARGOS_306 chromosome 2, complete sequence | 100 | 3.42E-139 | KWW26465.1 hypothetical protein AUK64_2547 [bacterium P201] | 61 | 5.90E-16 |
| k141_79388 | 296 | 121740.28 | CP022041 Prevotella melaninogenica strain FDAARGOS_306 chromosome 2, complete sequence | 99.662 | 2.86E-150 | EFC67102.1 LOW QUALITY PROTEIN: hypothetical protein HMPREF0670_02906, partial [Prevotella sp. oral taxon 317 str. F0108] | 84.2 | 3.00E-26 |
| **k141_275316** | **30474** | **120396** | **MG772933 Bat SARS-like coronavirus isolate bat-SL-CoVZC45, complete genome** | **89.113** | **0** | **AVP78030.1 non-structural polyprotein 1ab [Bat SARS-like coronavirus]** | **88.9** | **0** |
| k141_309125 | 412 | 107878.5 | CP022041 Prevotella melaninogenica strain FDAARGOS_306 chromosome 2, complete sequence | 98.058 | 0 | CUO89910.1 Uncharacterised protein [Prevotella copri] | 91.4 | 1.80E-21 |
| k141_79577 | 240 | 106402.31 | LC359515 Uncultured bacterium 83MF12012 gene for 16S rRNA, partial sequence | 100 | 6.52E-121 | OUD19662.1 hypothetical protein BUN10_26130 [Vibrio parahaemolyticus] | 82.7 | 2.20E-14 |
| k141_197704 | 409 | 104404.73 | CP023864 Prevotella jejuni strain CD3:33 chromosome II, complete sequence | 97.311 | 0 | KWW24027.1 hypothetical protein F082_2040 [bacterium F082] | 64.6 | 1.30E-11 |
| k141_280265 | 245 | 103339.01 | CP022041 Prevotella melaninogenica strain FDAARGOS_306 chromosome 2, complete sequence | 92.713 | 1.16E-93 | KWW26465.1 hypothetical protein AUK64_2547 [bacterium P201] | 61.7 | 1.20E-17 |
| k141_172622 | 290 | 100852.57 | CP023863 Prevotella jejuni strain CD3:33 chromosome I, complete sequence | 100 | 1.30E-148 | OPG95628.1 hypothetical protein B2I21_25150 [Paenibacillus sp. VT-16-81] | 72.9 | 7.80E-19 |
| k141_304224 | 301 | 99776.09 | CP023863 Prevotella jejuni strain CD3:33 chromosome I, complete sequence | 100 | 1.04E-154 | KWW24027.1 hypothetical protein F082_2040 [bacterium F082] | 66.7 | 7.40E-12 |
| k141_209219 | 671 | 99527.82 | CP019721 Veillonella parvula strain UTDB1-3, complete genome | 98.958 | 0 | EFG22293.1 hypothetical protein HMPREF0873_01746, partial [Veillonella sp. 3_1_44] | 98 | 2.60E-17 |
| k141_67655 | 719 | 95201.85 | CP023863 Prevotella jejuni strain CD3:33 chromosome I, complete sequence | 92.094 | 0 | WP_044045810.1 hypothetical protein [Prevotella melaninogenica] | 81.7 | 8.70E-27 |
| k141_205250 | 341 | 88773.84 | KF113907 Uncultured Prevotella sp. clone NA37_11 16S ribosomal RNA gene, partial sequence | 97.256 | 1.19E-154 | OXM99333.1 peptide YY, partial [Bifidobacterium vansinderenii] | 51.5 | 1.20E-13 |
| k141_179411 | 2733 | 87452.77 | CP012410 Leptotrichia sp. oral taxon 212 strain W10393, complete genome | 99.341 | 0 | EEX74022.1 hypothetical protein GCWU000323_01829 [Leptotrichia hofstadii F0254] | 85.6 | 2.10E-52 |
| k141_290049 | 476 | 85912.6 | CP013195 Prevotella enoeca strain F0113, complete genome | 95.607 | 0 | ETD26335.1 hypothetical protein HMPREF1173_02303 [Prevotella nigrescens CC14M] | 86.8 | 5.50E-70 |
| k141_51025 | 427 | 85350.28 | CP016205 Prevotella scopos JCM 17725 strain W2052 chromosome 2 genome | 99.766 | 0 | EFC67102.1 LOW QUALITY PROTEIN: hypothetical protein HMPREF0670_02906, partial [Prevotella sp. oral taxon 317 str. F0108] | 77 | 9.10E-32 |
| k141_46210 | 338 | 85072.61 | CP022041 Prevotella melaninogenica strain FDAARGOS_306 chromosome 2, complete sequence | 99.704 | 1.48E-173 | KWW26465.1 hypothetical protein AUK64_2547 [bacterium P201] | 73.1 | 5.30E-27 |

[a] Contig abundance evaluated as the excepted read counts by the RSEM program. For a transcript, the RSEM's expected counts may be slightly lower than the raw read counts due to the reads that map to multiple transcripts were divided among these transcripts.

**Table S2.** The top 80 abundant assembled contigs generated using the Trinity program.

| Contigs | Length | Abundance [a] | Result of blast against Nt database | | | Result of blast against Nr database | | |
|---|---|---|---|---|---|---|---|---|
| | | | Blast hit | Identity (%) | e-value | Blastx hit | Identity (%) | e-value |
| yingji_DN483576_c40_g3_i2 | 1923 | 2094180.66 | CP023863 Prevotella jejuni strain CD3:33 chromosome I, complete sequence | 97.558 | 0 | ETD26335.1 hypothetical protein HMPREF1173_02303 [Prevotella nigrescens CC14M] | 90.1 | 8.80E-66 |
| yingji_DN482282_c7_g3_i1 | 1426 | 1508548.65 | LT906445 Veillonella parvula strain NCTC11810 genome assembly, chromosome: 1 | 99.79 | 0 | ETJ17454.1 hypothetical protein Q620_VSAC00705G0001, partial [Veillonella sp. DORA_A_3_16_22] | 94.9 | 4.10E-76 |
| yingji_DN483576_c40_g3_i4 | 2115 | 957405.85 | CP022041 Prevotella melaninogenica strain FDAARGOS_306 chromosome 2, complete sequence | 90.297 | 0 | CDB46314.1 putative uncharacterized protein [Phascolarctobacterium sp. CAG:207] | 79.5 | 7.90E-84 |
| yingji_DN483576_c40_g1_i5 | 1227 | 747406.46 | AP018050 Prevotella melaninogenica DNA, complete genome, strain: GAI 07411 | 94.652 | 0 | KWW26465.1 hypothetical protein AUK64_2547 [bacterium P201] | 66.7 | 1.30E-25 |
| yingji_DN477344_c32_g1_i6 | 217 | 355553.84 | CP022040 Prevotella melaninogenica strain FDAARGOS_306 chromosome 1, complete sequence | 99.539 | 1.65E-106 | EHG15578.1 hypothetical protein HMPREF9138_01799, partial [Prevotella histicola F0411] | 97.8 | 6.10E-16 |
| yingji_DN483576_c40_g2_i1 | 793 | 353112.99 | GQ131418 Prevotella veroralis strain F0319 16S ribosomal RNA gene, partial sequence | 98.907 | 0 | EDM19152.1 hypothetical protein BACCAC_03786 [Bacteroides caccae ATCC 43185] | 78.6 | 7.80E-53 |
| yingji_DN482458_c5_g1_i13 | 275 | 343055.19 | AP018050 Prevotella melaninogenica DNA, complete genome, strain: GAI 07411 | 94.224 | 1.28E-113 | ETD26335.1 hypothetical protein HMPREF1173_02303 [Prevotella nigrescens CC14M] | 89.1 | 1.80E-25 |
| yingji_DN480761_c4_g1_i2 | 315 | 287176.88 | DQ537679 Uncultured bacterium clone B288-74 16S ribosomal RNA gene, partial sequence | 97.444 | 5.11E-148 | OPG95628.1 hypothetical protein B2I21_25150 [Paenibacillus sp. VT-16-81] | 69.5 | 4.50E-12 |
| yingji_DN481253_c3_g1_i4 | 234 | 281063.31 | LT678906 Prevotella melaninogenica partial 16S rRNA gene, isolate W538N_4320 | 99.134 | 1.38E-112 | OPG95628.1 hypothetical protein B2I21_25150 [Paenibacillus sp. VT-16-81] | 72 | 9.20E-10 |
| yingji_DN477344_c32_g1_i4 | 261 | 247610.82 | FJ557623 Uncultured bacterium clone ET_F_2c09 16S ribosomal RNA gene, partial sequence | 98.462 | 2.57E-125 | AOE06246.1 hypothetical protein [uncultured bacterium] | 60.9 | 1.10E-14 |
| yingji_DN482732_c3_g1_i1 | 353 | 224701.56 | CP023864 Prevotella jejuni strain CD3:33 chromosome II, complete sequence | 97.209 | 2.89E-96 | ETD26335.1 hypothetical protein HMPREF1173_02303 [Prevotella nigrescens CC14M] | 90.3 | 7.50E-24 |
| yingji_DN477344_c32_g2_i2 | 235 | 196262.23 | KY386203 Uncultured Prevotella sp. clone FAA299 16S ribosomal RNA gene, partial sequence | 99.574 | 1.78E-116 | OUD19662.1 hypothetical protein BUN10_26130 [Vibrio parahaemolyticus] | 83.3 | 1.80E-18 |
| yingji_DN474690_c4_g1_i4 | 556 | 192919.06 | CP002589 Prevotella denticola F0289, complete genome | 90.991 | 0 | KWW26465.1 hypothetical protein AUK64_2547 [bacterium P201] | 63.8 | 3.10E-32 |
| yingji_DN474759_c0_g1_i1 | 355 | 179075.38 | CP022040 Prevotella melaninogenica strain FDAARGOS_306 chromosome 1, complete sequence | 96.275 | 1.24E-159 | AOE06246.1 hypothetical protein [uncultured bacterium] | 57.9 | 1.10E-19 |
| yingji_DN482113_c3_g1_i2 | 222 | 155319.78 | KF113907 Uncultured Prevotella sp. clone NA37_11 16S ribosomal RNA gene, partial sequence | 97.748 | 1.32E-102 | EXT36960.1 hypothetical protein J810_4084, partial [Acinetobacter sp. 25977_7] | 73.2 | 1.30E-13 |
| yingji_DN481794_c4_g2_i1 | 329 | 150920.78 | LC356755 Uncultured bacterium 23MH11015 gene for 16S rRNA, partial sequence | 96.285 | 6.93E-147 | OUD19662.1 hypothetical protein BUN10_26130 [Vibrio parahaemolyticus] | 80.3 | 5.00E-22 |
| yingji_DN477344_c32_g3_i2 | 234 | 147867.65 | LT677940 Prevotella melaninogenica partial 16S rRNA gene, isolate 219N_3354 | 97.436 | 1.39E-107 | KMV77917.1 hypothetical protein HMPREF0979_01154, partial [Coprobacillus sp. 8_1_38FAA] | 75.7 | 5.60E-07 |
| yingji_DN482496_c5_g1_i9 | 230 | 147449.2 | CP023864 Prevotella jejuni strain CD3:33 chromosome II, complete sequence | 99.558 | 1.75E-111 | EFN91701.1 hypothetical protein HMPREF9018_1166 [Prevotella amnii CRIS 21A-A] | 96 | 1.40E-31 |
| yingji_DN479682_c3_g1_i23 | 253 | 145114.61 | KY386203 Uncultured Prevotella sp. clone FAA299 16S ribosomal RNA gene, partial sequence | 98.814 | 4.14E-123 | OUD19662.1 hypothetical protein BUN10_26130 [Vibrio parahaemolyticus] | 78.8 | 3.10E-19 |
| yingji_DN474567_c2_g4_i1 | 242 | 142374.1 | CP023863 Prevotella jejuni strain CD3:33 chromosome I, complete sequence | 95.816 | 1.13E-103 | KDS36881.1 hypothetical protein M091_0855 [Parabacteroides distasonis str. 3776 D15 i] | 67.6 | 2.20E-14 |
| yingji_DN475086_c3_g1_i12 | 601 | 134057.33 | CP023864 Prevotella jejuni strain CD3:33 chromosome II, complete sequence | 96.179 | 0 | KWW24027.1 hypothetical protein F082_2040 [bacterium F082] | 60.9 | 2.10E-26 |
| yingji_DN483576_c40_g2_i2 | 518 | 129481.81 | KP294789Uncultured Veillonella sp. clone P17-29-T7 16S ribosomal RNA gene, partial sequence | 95.402 | 0 | EDP22130.1 hypothetical protein FAEPRAM212_01166 [Faecalibacterium prausnitzii M21/2] | 78.4 | 5.30E-50 |
| yingji_DN477344_c32_g1_i7 | 424 | 128501.09 | CP023863 Prevotella jejuni strain CD3:33 chromosome I, complete sequence | 97.866 | 2.51E-157 | AOE06246.1 hypothetical protein [uncultured bacterium] | 61.3 | 2.00E-23 |
| yingji_DN481203_c1_g1_i8 | 330 | 119489.42 | CP023863 Prevotella jejuni strain CD3:33 chromosome I, complete sequence | 97.77 | 7.16E-127 | EFC67102.1 LOW QUALITY PROTEIN: hypothetical protein HMPREF0670_02906, partial [Prevotella sp. oral taxon 317 str. F0108] | 75 | 3.90E-30 |
| yingji_DN482113_c3_g1_i10 | 222 | 117885.78 | GQ398420 Uncultured bacterium clone 47 16S ribosomal RNA gene, partial sequence | 97.596 | 7.98E-95 | EXT36960.1 hypothetical protein J810_4084, partial [Acinetobacter sp. 25977_7] | 66.1 | 8.70E-10 |
| yingji_DN477344_c32_g1_i1 | 412 | 117747.68 | FJ557623 Uncultured bacterium clone ET_F_2c09 16S ribosomal RNA gene, partial sequence | 98.403 | 6.82E-153 | AOE06246.1 hypothetical protein [uncultured bacterium] | 60.7 | 4.40E-23 |
| yingji_DN477518_c5_g2_i5 | 227 | 117518.07 | CP023863 Prevotella jejuni strain CD3:33 chromosome I, complete sequence | 96.847 | 2.93E-99 | AOE11741.1 hypothetical protein [uncultured bacterium] | 67.2 | 1.30E-13 |

| | | | | | | | | |
|---|---|---|---|---|---|---|---|---|
| yingji_DN482535_c5_g1_i14 | 299 | 107128.47 | JF123172 Uncultured bacterium clone ncd1418b06c1 16S ribosomal RNA gene, partial sequence | 99.663 | 8.03E-151 | ABZ84906.1 hypothetical protein HM1_3148 [Heliobacterium modesticaldum Ice1] | 77.3 | 2.10E-27 |
| yingji_DN482535_c5_g1_i4 | 237 | 106872.4 | LT906445 Veillonella parvula strain NCTC11810 genome assembly, chromosome: 1 | 100 | 4.15E-68 | CRE39519.1 transposase for IS1272 [Staphylococcus aureus] | 70.6 | 1.40E-10 |
| yingji_DN478259_c6_g1_i2 | 227 | 96776.77 | CP023863 Prevotella jejuni strain CD3:33 chromosome I, complete sequence | 94.416 | 1.40E-77 | CUO90010.1 Uncharacterised protein [Prevotella copri] | 90.6 | 4.10E-07 |
| yingji_DN474690_c4_g1_i8 | 314 | 90964.48 | CP016205 Prevotella scopos JCM 17725 strain W2052 chromosome 2 genome | 92.089 | 1.14E-119 | KWW26465.1 hypothetical protein AUK64_2547 [bacterium P201] | 60.4 | 6.30E-22 |
| yingji_DN479388_c0_g1_i2 | 265 | 88879.77 | LC358495 Uncultured bacterium 62MG02014 gene for 16S rRNA, partial sequence | 95.802 | 9.52E-115 | OUD19662.1 hypothetical protein BUN10_26130 [Vibrio parahaemolyticus] | 77.1 | 1.60E-18 |
| yingji_DN482535_c5_g1_i5 | 220 | 85474.58 | DQ394709 Veillonella parvula strain H2 16S ribosomal RNA gene, partial sequence | 99.091 | 1.67E-106 | CDN41090.1 hypothetical protein BN871_AB_00880 [Paenibacillus sp. P22] | 67.1 | 3.10E-15 |
| yingji_DN483576_c40_g2_i4 | 794 | 85025.94 | JQ460268 Uncultured bacterium clone 070054_517 16S ribosomal RNA gene, partial sequence | 96.343 | 0 | EDP22130.1 hypothetical protein FAEPRAM212_01166 [Faecalibacterium prausnitzii M21/2] | 66.9 | 2.40E-49 |
| yingji_DN483239_c4_g4_i1 | 379 | 80255.5 | CP023864 Prevotella jejuni strain CD3:33 chromosome II, complete sequence | 96.477 | 1.01E-170 | EFN91701.1 hypothetical protein HMPREF9018_1166 [Prevotella amnii CRIS 21A-A] | 87.9 | 5.60E-49 |
| yingji_DN475296_c6_g1_i10 | 414 | 76992.72 | JQ448356 Uncultured bacterium clone 069077_255 16S ribosomal RNA gene, partial sequence | 95.844 | 0 | AOE06246.1 hypothetical protein [uncultured bacterium] | 60 | 2.70E-20 |
| yingji_DN480267_c2_g1_i4 | 210 | 75430.95 | JQ077772 Uncultured bacterium clone HAV7D9G02BX98V 16S ribosomal RNA gene, partial sequence | 99.383 | 5.94E-76 | OUD19662.1 hypothetical protein BUN10_26130 [Vibrio parahaemolyticus] | 82.4 | 1.50E-14 |
| yingji_DN475296_c6_g1_i2 | 476 | 75075.24 | AM420082 Uncultured Prevotella sp. partial 16S rRNA gene, clone 302B04(oral) | 96.603 | 0 | AOE06246.1 hypothetical protein [uncultured bacterium] | 62.9 | 7.80E-24 |
| yingji_DN481323_c5_g2_i2 | 278 | 74663.27 | CP023863 Prevotella jejuni strain CD3:33 chromosome I, complete sequence | 91.786 | 1.32E-103 | KWW26465.1 hypothetical protein AUK64_2547 [bacterium P201] | 56 | 9.80E-19 |
| yingji_DN480267_c2_g1_i2 | 308 | 73945.05 | JQ471950 Uncultured bacterium clone 071054_096 16S ribosomal RNA gene, partial sequence | 96.644 | 2.35E-136 | OUD19662.1 hypothetical protein BUN10_26130 [Vibrio parahaemolyticus] | 82.4 | 2.10E-14 |
| yingji_DN480509_c0_g1_i1 | 257 | 72308.97 | LT684910 Uncultured Prevotella sp. partial 16S rRNA gene, isolate W787N_10325 | 96.996 | 2.59E-105 | OUD19662.1 hypothetical protein BUN10_26130 [Vibrio parahaemolyticus] | 73.4 | 8.50E-17 |
| yingji_DN479926_c4_g1_i4 | 222 | 70429.24 | MH078430 Uncultured Capnocytophaga sp. clone 174_p8_c_25 16S ribosomal RNA gene, partial sequence | 98.013 | 1.79E-66 | AOE12499.1 hypothetical protein [uncultured bacterium] | 68 | 3.40E-06 |
| yingji_DN479135_c8_g1_i2 | 264 | 70245.75 | CP022041 Prevotella melaninogenica strain FDAARGOS_306 chromosome 2, complete sequence | 98.333 | 8.04E-51 | CUO90010.1 Uncharacterised protein [Prevotella copri] | 93.8 | 5.70E-08 |
| yingji_DN482458_c5_g1_i11 | 292 | 67762.99 | CP022041 Prevotella melaninogenica strain FDAARGOS_306 chromosome 2, complete sequence | 95.848 | 4.84E-128 | ETD26335.1 hypothetical protein HMPREF1173_02303 [Prevotella nigrescens CC14M] | 86.7 | 1.30E-18 |
| yingji_DN483275_c3_g1_i23 | 321 | 64351.2 | CP023864 Prevotella jejuni strain CD3:33 chromosome II, complete sequence | 95.912 | 1.89E-142 | EFN91701.1 hypothetical protein HMPREF9018_1166 [Prevotella amnii CRIS 21A-A] | 85.9 | 2.80E-33 |
| yingji_DN477344_c31_g1_i2 | 388 | 61080.74 | LT679278 Prevotella melaninogenica partial 16S rRNA gene, isolate 43T_4692 | 96.392 | 1.32E-179 | OPG95628.1 hypothetical protein B2I21_25150 [Paenibacillus sp. VT-16-81] | 59 | 2.60E-17 |
| yingji_DN474678_c1_g1_i10 | 641 | 58815.68 | JQ478347 Uncultured bacterium clone 071076_162 16S ribosomal RNA gene, partial sequence | 94.543 | 0 | KFJ04251.1 PG1 protein [Bifidobacterium thermacidophilum subsp. thermacidophilum] | 50.8 | 1.70E-18 |
| yingji_DN477344_c33_g1_i6 | 201 | 58581.35 | EU993256 Uncultured bacterium clone WG_e55 16S ribosomal RNA gene, partial sequence | 95.522 | 2.59E-84 | | | |
| yingji_DN470028_c1_g1_i1 | 314 | 58418.12 | JQ470050 Uncultured bacterium clone 071024_066 16S ribosomal RNA gene, partial sequence | 93.98 | 1.47E-123 | EXT36960.1 hypothetical protein J810_4084, partial [Acinetobacter sp. 25977_7] | 66.1 | 1.00E-08 |
| yingji_DN474678_c1_g1_i3 | 366 | 57874.08 | LT677940 Prevotella melaninogenica partial 16S rRNA gene, isolate 219N_3354 | 98.361 | 0 | OPG95628.1 hypothetical protein B2I21_25150 [Paenibacillus sp. VT-16-81] | 68 | 3.40E-19 |
| yingji_DN482282_c7_g1_i5 | 295 | 55295.31 | CP019721 Veillonella parvula strain UTDB1-3, complete genome | 97.288 | 4.83E-138 | ABZ84885.1 hypothetical protein HM1_3125 [Heliobacterium modesticaldum Ice1] | 54.2 | 7.70E-06 |
| yingji_DN477344_c32_g3_i1 | 251 | 54781.57 | LT688914 Prevotella nanceiensis partial 16S rRNA gene, isolate W840T_14330 | 97.61 | 5.35E-117 | EXY63944.1 hypothetical protein M085_3631 [Bacteroides fragilis str. 3986 N(B)19] | 60.9 | 9.80E-10 |
| yingji_DN477344_c32_g2_i4 | 371 | 54773.47 | GQ365015 Uncultured bacterium clone 89BAL_G12 16S ribosomal RNA gene, partial sequence | 98.638 | 0 | EDO51672.1 hypothetical protein BACUNI_04219 [Bacteroides uniformis ATCC 8492] | 71.3 | 1.80E-23 |
| yingji_DN480761_c4_g1_i1 | 205 | 51418.06 | LT688896 Prevotella melaninogenica partial 16S rRNA gene, isolate W840T_14312 | 97.537 | 4.37E-92 | OPG95628.1 hypothetical protein B2I21_25150 [Paenibacillus sp. VT-16-81] | 63.9 | 9.50E-11 |
| yingji_DN483048_c4_g1_i4 | 233 | 49367.6 | CP022041 Prevotella melaninogenica strain FDAARGOS_306 chromosome 2, complete sequence | 96.957 | 1.08E-103 | KWW25567.1 Uncharacterized protein AUK64_2610, partial [bacterium P201] | 84.8 | 3.90E-13 |
| yingji_DN482282_c7_g1_i2 | 405 | 48841.63 | CP019721 Veillonella parvula strain UTDB1-3, complete genome | 98.765 | 0 | CUP36263.1 Uncharacterised protein [Bacteroides xylanisolvens] | 63 | 1.30E-16 |
| yingji_DN483576_c39_g1_i2 | 384 | 48687.59 | CP023863 Prevotella jejuni strain CD3:33 chromosome I, complete sequence | 94.531 | 1.03E-165 | KWW26465.1 hypothetical protein AUK64_2547 [bacterium P201] | 56.8 | 8.20E-24 |
| yingji_DN479496_c2_g1_i1 | 373 | 46219.95 | JN382502 Uncultured bacterium clone ZB1881012 16S ribosomal RNA gene, partial sequence | 96.196 | 5.99E-168 | AOE11686.1 hypothetical protein [uncultured bacterium] | 68.2 | 1.20E-19 |
| yingji_DN476234_c3_g1_i1 | 269 | 46071.94 | LC356684 Uncultured bacterium 23MB01003 gene for 16S rRNA, partial sequence | 95.911 | 1.24E-118 | OUD19662.1 hypothetical protein BUN10_26130 [Vibrio parahaemolyticus] | 74.3 | 4.20E-19 |

| | | | | | | | | |
|---|---|---|---|---|---|---|---|---|
| yingji_DN481203_c1_g1_i21 | 248 | 45956.1 | CP023864 Prevotella jejuni strain CD3:33 chromosome II, complete sequence | 97.177 | 1.14E-113 | EFC67102.1 LOW QUALITY PROTEIN: hypothetical protein HMPREF0670_02906, partial [Prevotella sp. oral taxon 317 str. F0108] | 82.4 | 2.90E-22 |
| yingji_DN483576_c40_g3_i9 | 293 | 45171.15 | CP003667 Prevotella sp. oral taxon 299 str. F0039 plasmid, complete sequence | 99.317 | 6.11E-147 | EFN91701.1 hypothetical protein HMPREF9018_1166 [Prevotella amnii CRIS 21A-A] | 96.9 | 3.10E-47 |
| yingji_DN481203_c1_g1_i12 | 306 | 43194.56 | CP022041 Prevotella melaninogenica strain FDAARGOS_306 chromosome 2, complete sequence | 97.712 | 8.27E-146 | EFC67102.1 LOW QUALITY PROTEIN: hypothetical protein HMPREF0670_02906, partial [Prevotella sp. oral taxon 317 str. F0108] | 80.9 | 1.00E-21 |
| yingji_DN478175_c2_g1_i2 | 571 | 42741.6 | CP016205 Prevotella scopos JCM 17725 strain W2052 chromosome 2 genome | 90.698 | 0 | KWW26465.1 hypothetical protein AUK64_2547 [bacterium P201] | 75 | 8.70E-30 |
| yingji_DN482458_c5_g1_i7 | 248 | 41176.22 | CP022041 Prevotella melaninogenica strain FDAARGOS_306 chromosome 2, complete sequence | 98.367 | 5.27E-117 | EDY97039.1 hypothetical protein BACPLE_00421 [Bacteroides plebeius DSM 17135] | 80.3 | 2.60E-23 |
| yingji_DN479926_c1_g1_i1 | 496 | 40272.89 | FM997688 Uncultured bacterium partial 16S rRNA gene, clone 16sps27-5a05.w2k | 95.749 | 0 | OPG95628.1 hypothetical protein B2I21_25150 [Paenibacillus sp. VT-16-81] | 58.1 | 1.40E-12 |
| yingji_DN482529_c2_g1_i1 | 473 | 39126.62 | CP022041 Prevotella melaninogenica strain FDAARGOS_306 chromosome 2, complete sequence | 93.137 | 1.00E-166 | EFN91701.1 hypothetical protein HMPREF9018_1166 [Prevotella amnii CRIS 21A-A] | 83.7 | 4.10E-33 |
| yingji_DN483576_c40_g3_i3 | 208 | 38527.64 | CP023864 Prevotella jejuni strain CD3:33 chromosome II, complete sequence | 98.558 | 3.41E-98 | CUO89876.1 Cell wall-associated hydrolase [Prevotella copri] | 95.7 | 7.80E-29 |
| yingji_DN482113_c2_g1_i4 | 354 | 37700.68 | CP019721 Veillonella parvula strain UTDB1-3, complete genome | 98.58 | 1.21E-174 | EFG22293.1 hypothetical protein HMPREF0873_01746, partial [Veillonella sp. 3_1_44] | 97.7 | 2.40E-14 |
| yingji_DN483110_c4_g1_i10 | 240 | 35521.85 | EU063557 Uncultured bacterium clone LM0ACA28ZD06FM1 genomic sequence | 86.364 | 1.18E-63 | EFU29156.1 hypothetical protein HMPREF6485_2897, partial [Prevotella buccae ATCC 33574] | 48.5 | 4.20E-10 |
| yingji_DN469226_c0_g1_i1 | 325 | 34281.13 | CP023863 Prevotella jejuni strain CD3:33 chromosome I, complete sequence | 94.044 | 5.40E-133 | AOE06246.1 hypothetical protein [uncultured bacterium] | 55.2 | 7.40E-18 |
| yingji_DN483110_c4_g1_i5 | 279 | 33609.11 | AP018050 Prevotella melaninogenica DNA, complete genome, strain: GAI 07411 | 94.203 | 4.70E-113 | KDS36881.1 hypothetical protein M091_0855 [Parabacteroides distasonis str. 3776 D15 i] | 67.1 | 1.80E-20 |
| yingji_DN480296_c1_g4_i1 | 202 | 33474.87 | CP003667 Prevotella sp. oral taxon 299 str. F0039 plasmid, complete sequence | 97.525 | 1.54E-91 | EDO14276.1 hypothetical protein BACOVA_00014 [Bacteroides ovatus ATCC 8483] | 57.6 | 2.30E-09 |
| yingji_DN474678_c1_g1_i1 | 211 | 33410.85 | LT677940 Prevotella melaninogenica partial 16S rRNA gene, isolate 219N_3354 | 97.63 | 1.62E-96 | AOE06246.1 hypothetical protein [uncultured bacterium] | 50.7 | 1.90E-06 |
| **yingji_DN483566_c8_g3_i1** | **11760** | **33252** | **MG772933 Bat SARS-like coronavirus isolate bat-SL-CoVZC45, complete genome** | **90.415** | **0** | **AVP78030.1 non-structural polyprotein 1ab [Bat SARS-like coronavirus]** | **97.3** | **0** |
| yingji_DN482535_c5_g1_i2 | 214 | 33075.64 | JQ457132 Uncultured bacterium clone 070007_385 16S ribosomal RNA gene, partial sequence | 98.095 | 1.27E-97 | OBZ15173.1 hypothetical protein A7975_32355 [Bacillus sp. FJAT-26390] | 73.3 | 8.60E-15 |
| yingji_DN482627_c4_g1_i6 | 848 | 32710.61 | CP012072 Actinomyces meyeri strain W712, complete genome | 89.711 | 0 | KMS64810.1 hypothetical protein BVRB_042430, partial [Beta vulgaris subsp. vulgaris] | 63.6 | 1.30E-24 |
| yingji_DN481441_c5_g1_i8 | 369 | 32150.86 | CP016205 Prevotella scopos JCM 17725 strain W2052 chromosome 2 genome | 97.561 | 2.10E-177 | KWW26465.1 hypothetical protein AUK64_2547 [bacterium P201] | 71 | 6.50E-26 |
| yingji_DN481434_c1_g1_i7 | 562 | 31243.87 | CP012072 Actinomyces meyeri strain W712, complete genome | 92.568 | 1.19E-176 | EDX25829.1 conserved hypothetical protein [Streptomyces sp. Mg1] | 53.1 | 1.50E-26 |
| yingji_DN476965_c6_g1_i1 | 344 | 30550.91 | CP024735 Prevotella intermedia strain KCOM 1944 chromosome 2, complete sequence | 94.671 | 2.66E-136 | KWW26465.1 hypothetical protein AUK64_2547 [bacterium P201] | 82.1 | 1.70E-17 |
| yingji_DN481203_c1_g1_i19 | 379 | 30484.35 | CP022041 Prevotella melaninogenica strain FDAARGOS_306 chromosome 2, complete sequence | 97.098 | 1.29E-179 | KWW26465.1 hypothetical protein AUK64_2547 [bacterium P201] | 66.2 | 5.10E-18 |

[a] Contig abundance was evaluated as the excepted read counts by the RSEM program. For a transcript, the RSEM's expected count may be slightly lower than the raw read count due to the reads that map to multiple transcripts were divided among these transcripts.

**Table S3.** Amino acid identities of the selected predicted gene products between the novel coronavirus (WHCV) and known betacoronaviruses.

| CoV | Strains | Amino acid identity (%) | | | | | | | | | | | | | | | | | | | | | |
|---|---|---|---|---|---|---|---|---|---|---|---|---|---|---|---|---|---|---|---|---|---|---|---|
| | | nsp1 | nsp2 | nsp3 | nsp4 | nsp5 | nsp6 | nsp7 | nsp8 | nsp9 | nsp10 | nsp11 | nsp12 | nsp13 | nsp14 | nsp15 | nsp16 | S | ORF3 | E | M | ORF8 | N |
| *Sarbecovirus* | Bat-SL-CoVZC45 | 84.4 | 95.3 | 94.4 | 96.8 | 99.0 | 97.9 | 100 | 97.5 | 97.3 | 97.1 | 85.7 | 95.9 | 99.3 | 94.5 | 89.0 | 98.0 | 82.3 | 90.9 | 100 | 98.7 | 94.3 | 94.0 |
| | SARS-CoV Tor2 | 95.6 | 68.3 | 77.3 | 79.9 | 51.2 | 87.2 | 98.8 | 97.5 | 97.3 | 97.1 | 85.7 | 96.3 | 99.8 | 95.1 | 88.7 | 93.3 | 77.2 | 72.7 | 96.1 | 91.0 | 28.0 | 91.0 |
| | BM48-31/BGR/2008 | 81.7 | 62.5 | 72.9 | 81.1 | 94.1 | 83.8 | 95.2 | 96.5 | 98.2 | 94.3 | 78.6 | 95.4 | 97.8 | 93.5 | 89.9 | 88.6 | 73.2 | 63.6 | 93.4 | 87.9 | / | 88.2 |
| | WIV1 | 85.0 | 67.3 | 77.0 | 80.3 | 95.8 | 86.9 | 100 | 97.5 | 97.3 | 97.9 | 85.7 | 96.4 | 99.5 | 95.4 | 89.0 | 93.0 | 78.3 | 74.5 | 96.1 | 90.1 | 58.2 | 91.0 |
| | JTMC15 | 78.9 | 68.9 | 76.0 | 81.3 | 94.8 | 85.9 | 98.8 | 96.5 | 97.3 | 97.1 | 85.7 | 96.4 | 98.5 | 94.9 | 88.2 | 92.6 | 74.3 | 68.4 | 92.1 | 90.5 | / | 90.0 |
| *Merbecovirus* | EriCoV | 16.5 | 18.6 | 30.1 | 42.4 | 49.2 | 34.6 | 60.2 | 52.8 | 50.0 | 60.7 | 46.2 | 71.3 | 71.1 | 63.6 | 50.0 | 65.8 | 29.3 | / | 40.8 | 43.1 | / | 46.8 |
| | Ty-BatCoV-HKU4 | 17.0 | 17.5 | 30.6 | 37.2 | 68.6 | 34.9 | 54.2 | 50.8 | 51.8 | 59.0 | 53.8 | 70.8 | 70.9 | 63.0 | 50.9 | 65.4 | 31.7 | / | 40.8 | 42.0 | / | 47.0 |
| | MERS-CoV | 16.6 | 18.3 | 30.1 | 39.2 | 50.8 | 33.9 | 55.4 | 52.8 | 53.6 | 58.6 | 46.2 | 71.3 | 71.6 | 63.6 | 50.9 | 66.1 | 27.4 | / | 35.5 | 40.6 | / | 47.0 |
| | Pi-BatCoV_HKU5 | 17.5 | 18.1 | 30.6 | 39.8 | 52.3 | 34.3 | 56.6 | 51.3 | 48.2 | 56.1 | 46.2 | 71.6 | 71.7 | 62.8 | 51.5 | 65.4 | 27.5 | / | 32.9 | 41.6 | / | 45.2 |
| *Nobecovirus* | Ro-BatCoV_GCCDC1 | 24.1 | 16.2 | 29.4 | 40.9 | 52.0 | 36.2 | 66.3 | 57.6 | 55.4 | 62.9 | 38.5 | 72.3 | 73.7 | 61.6 | 49.7 | 63.5 | 32.1 | / | 32.4 | 43.4 | / | 39.7 |
| | Ro-BatCoV_HKU9 | 26.4 | 19.1 | 30.4 | 43.1 | 50.2 | 33.6 | 67.5 | 57.6 | 58 | 65.0 | 38.5 | 72.6 | 74.0 | 61.2 | 47.5 | 62.3 | 31.4 | / | 28.4 | 39.6 | / | 41.5 |
| *Hibecovirus* | Bat_Hp-BetaCoV | 23.8 | 27.0 | 38.6 | 53.8 | 49.2 | 44.8 | 72.3 | 60.1 | 61.6 | 68.6 | 69.2 | 77.5 | 80.7 | 70.2 | 61.6 | 67.8 | 42.8 | / | 53.9 | 52.5 | / | 50.1 |
| *Embecovirus* | HCoV_HKU1 | 12.7 | 11.8 | 22.2 | 41.3 | 47.9 | 28.5 | 47.0 | 46.6 | 46.4 | 52.6 | 61.5 | 67.0 | 65.3 | 58.3 | 49.1 | 63.4 | 27.4 | / | 28.4 | 36.4 | / | 31.6 |
| | HCoV_OC43 | 15.2 | 10.8 | 22.8 | 41.5 | 50.8 | 28.5 | 49.4 | 47.4 | 46.4 | 51.8 | 61.5 | 65.4 | 67.9 | 58.0 | 47.6 | 66.1 | 28.4 | / | 22.4 | 40.1 | / | 32.0 |
| | ChRCoV_HKU24 | 16.5 | 11.3 | 22.3 | 40.4 | 99.0 | 29.9 | 48.2 | 46.4 | 44.5 | 54.0 | 61.5 | 67.0 | 68.8 | 59.2 | 48.8 | 65.1 | 27.8 | / | 25.0 | 37.2 | / | 31.1 |
| | MHV | 17.4 | 10.3 | 23.0 | 41.1 | 50.2 | 28.5 | 44.6 | 47.1 | 48.2 | 52.6 | 61.5 | 65.8 | 67.3 | 58.1 | 48.2 | 63.1 | 28.1 | / | 25.0 | 39.2 | / | 32.4 |

**Table S4.** Cleavage products of the replicase polyproteins of WHCV.

| Cleavage product | Position in polyprotein pp1a/pp1ab[a] | Protein size (no. of amino acids) | Putative functional domain(s)[b] |
|---|---|---|---|
| nsp1 | 1Met-Gly180 | 180 | |
| nsp2 | 181Ala-Gly818 | 638 | |
| nsp3 | 819Ala-Gly2763 | 1945 | ADRP |
| nsp4 | 2764Lys-Gln3263 | 500 | |
| nsp5 | 3264Ser-Gln3569 | 306 | 3CLpro |
| nsp6 | 3570Ser-Gln3859 | 290 | |
| nsp7 | 3860Ser-Gln3942 | 83 | |
| nsp8 | 3943Ala-Gln4140 | 198 | |
| nsp9 | 4141Asn-Gln4253 | 113 | |
| nsp10 | 4254Ala-Gln4392 | 139 | |
| nsp11 | 4393Ser-Val4405 | 13 | |
| nsp12 | 4393Ser-Gln5324 | 932 | RdRp |
| nsp13 | 5325Ala-Gln5925 | 601 | Hel |
| nsp14 | 5926Ala-Gln6452 | 527 | ExoN |
| nsp15 | 6453Ser-Gln6798 | 346 | NendoU |
| nsp16 | 6799Ser-Asn7096 | 298 | O-MT |

[a]Amino acids of replicase proteins pp1a and pp1ab were numbered with the assumption that a -1 ribosomal frameshift occurs to express ORF1b, and use of the slippery sequence UUUAAAC is predicted to yield a peptide bond between Asn4401 and Arg4402 in pp1ab.

[b]Abbreviations: ADRP, adenosine diphosphate-ribose 1”-phosphatase; 3CLpro, 3C-like cysteine proteinase; RdRp, RNA-dependent RNA polymerase; Hel, helicase; ExoN, 3’-to-5’ exonuclease; NendoU, nidoviral endoribonuclease specific for U; OMT, S-adenosylmethionine-dependent ribose 2’-O-methyltransferase.

**Table S5.** Predicted gene functions of WHCV ORFs.

| ORF name | Proposed function |
|---|---|
| ORF 1a | Encoded nonstructural proteins (nsp1 to nsp11), essential for viral replication, viral assembly, immune response modulation, etc. |
| ORF 1b | Encoded nonstructural proteins (nsp12 to nsp16), essential for viral replication |
| S | Spike protein, binding to cell receptor and mediate virus-cell fusion |
| ORF 3a | Accessory protein |
| ORF 3b | Accessory protein |
| E | Envelope protein, virus assembly and morphogenesis |
| M | Membrane protein, virus assembly |
| ORF6 | Accessory protein |
| ORF 7a | Accessory protein |
| ORF 7b | Accessory protein |
| ORF8 | Accessory protein |
| N | Nucleocapsid protein, forms complexes with genomic RNA, interact with M protein for viral assembly |
| ORF 9a | Accessory protein |
| ORF 9b | Accessory protein |
| ORF 10 | Accessory protein |

**Table S6**. Coding of potential and putative transcription regulatory sequences of the genome sequence of WHCV.

| ORF | Location (nt) | Length (nt) | Length (aa) | TRS location | TRS sequence (s) (distance in bases to AUG) |
|---|---|---|---|---|---|
| 1ab | 266-21,555 (shift at13,468) | 21,290 | 7,096 | 64 | CU**CUAAACGAAC**UU(188)[a]AUG |
| S | 21,563-25,384 | 3,822 | 1,273 | 21,550 | AA**CUAAACGAAC**AAUG |
| 3a | 25,393-26,220 | 828 | 275 | 25,379 | ACAUAA**ACGAAC**UUAUG |
| 3b | 25,765-26,220 | 456 | 151 | | |
| E | 26,245-26,472 | 228 | 75 | 26,231 | AUGAGU**ACGAAC**UUAUG |
| M | 26,523-27,191 | 669 | 222 | 26,467 | GU**CUAAACGAAC**UA(42)[a]AUG |
| 6 | 27,202-27,387 | 186 | 61 | 27,035 | UACAUC**ACGAAC**GC(153)[a] AUG |
| 7a | 27,394-27,759 | 366 | 121 | 27,382 | GAUUAA**ACGAAC**AUG |
| 7b | 27,756-27,887 | 132 | 43 | | |
| 8 | 27,894-28,259 | 366 | 121 | 27,882 | GC**CUAAACGAAC**AUG |
| N | 28,274-29,533 | 1,260 | 419 | 28,254 | AU**CUAAACGAAC**AA(6)[a]AUG |
| 9a | 28,284-28,577 | 294 | 97 | | |
| 9b | 28,734-28,955 | 222 | 73 | | |
| 10 | 29,558-29,674 | 117 | 38 | 29,528 | GC**CUAAAC**UCAUGC(16)[a] AUG |

[a]Numbers in parentheses represent the number of nucleotides to the putative start codon. Start codons are underlined. The conserved TRS core sequence, ACGAAC or CUAAAC, is highlighted in bold.

**Table S7.** Amino acid identities of the RBD sequence between SARS- and bat SARS-like CoVs.

| | 1 | 2 | 3 | 4 | 5 | 6 | 7 | 8 | 9 | 10 | 11 | 12 | 13 |
|---|---|---|---|---|---|---|---|---|---|---|---|---|---|
| **1. SARS-CoV_Tor2** | | 100 | 100 | 97.9 | 96.4 | 95.4 | 80.9 | 73.8 | 62.4 | 62.4 | 62.4 | 61.9 | 62.9 |
| **2. SARS-CoV_BJ01** | 100 | | 100 | 97.9 | 96.4 | 95.4 | 80.9 | 73.8 | 62.4 | 62.4 | 62.4 | 61.9 | 62.9 |
| **3. SARS-CoV_WH20** | 100 | 100 | | 97.9 | 96.4 | 95.4 | 80.9 | 73.8 | 62.4 | 62.4 | 62.4 | 61.9 | 62.9 |
| **4. SARS-CoV_SZ3** | 97.9 | 97.9 | 97.9 | | 96.9 | 95.9 | 82.0 | 74.9 | 62.4 | 62.4 | 62.4 | 61.9 | 62.9 |
| **5. Bat_SL_Rs7327** | 96.4 | 96.4 | 96.4 | 96.9 | | 97.9 | 83.0 | 75.9 | 62.9 | 62.4 | 62.9 | 61.9 | 63.4 |
| **6.Bat_SL_Rs4874** | 95.4 | 95.4 | 95.4 | 95.9 | 97.9 | | 82.0 | 76.4 | 63.4 | 63.4 | 63.4 | 62.9 | 63.9 |
| **7.Bat_SL_Rs4231** | 80.9 | 80.9 | 80.9 | 82.0 | 83.0 | 82.0 | | 76.9 | 62.4 | 62.4 | 62.4 | 61.9 | 62.9 |
| **8. WH-human 1** | 73.8 | 73.8 | 73.8 | 74.9 | 75.9 | 76.4 | 76.9 | | 63.6 | 64.1 | 64.1 | 63.6 | 64.6 |
| **9. Bat_SL_CoVZC45** | 62.4 | 62.4 | 62.4 | 62.4 | 62.9 | 63.4 | 62.4 | 63.6 | | 91.5 | 95.5 | 88.6 | 91.0 |
| **10.Bat_SL_Rp3** | 62.4 | 62.4 | 62.4 | 62.4 | 62.4 | 63.4 | 62.4 | 64.1 | 91.5 | | 90.3 | 96.0 | 95.5 |
| **11. Bat_SL_Rf1** | 62.4 | 62.4 | 62.4 | 62.4 | 62.9 | 63.4 | 62.4 | 64.1 | 95.5 | 90.3 | | 88.1 | 89.3 |
| **12. Bat_SL_Rm1** | 61.9 | 61.9 | 61.9 | 61.9 | 61.9 | 62.9 | 61.9 | 63.6 | 88.6 | 96.0 | 88.1 | | 92.7 |
| **13. Bat_SL_HKU3** | 62.9 | 62.9 | 62.9 | 62.9 | 63.4 | 63.9 | 62.9 | 64.6 | 91.0 | 95.5 | 89.3 | 92.7 | |

**Table S8.** PCR primers used in this study.

| Primer name | Sequence (5`-3`) | Region/Size |
|---|---|---|
| A. Primers for entire genome amplification | | |
| WHCV-F1 | CCAGGTAACAAACCAACCAACTT | 36-58 |
| WHCV-R1 | GGCAACCAACATAAGAGAACACAC | 1507-1530 |
| WHCV-F2 | CAACCAAATGTGCCTTTCAACTC | 1217-1239 |
| WHCV-R2 | CACAGTGTCATCACCAAAAGTAACCT | 2746-2771 |
| WHCV-F3 | TGTCACGCACTCAAAGGGATT | 2408-2428 |
| WHCV-R3 | GACAGCTAAGTAGACATTTGTGCGAA | 3787-3812 |
| WHCV-F4 | ATGCCATGCAAGTTGAATCTGAT | 3523-3545 |
| WHCV-R4 | TGCGTGTGGAGGTTAATGTTGT | 5005-5026 |
| WHCV-F5 | GATCTCTCAAAGTGCCAGCTACAGT | 4681-4705 |
| WHCV-R5 | TTATAATCAATAGCCACCACATCACC | 6174-6199 |
| WHCV-F6 | AGAAACTTTGTATTGCATAGACGGTG | 5807-5832 |
| WHCV-R6 | ACCAGTACAGTAAGAAGGCATGCC | 7053-7076 |
| WHCV-F7 | GTTTAGCTGCTGTTAATAGTGTCCCTT | 6658-6684 |
| WHCV-R7 | TGCAACTTCCGCACTATCACC | 8022-8042 |
| WHCV-F8 | TCCTACTGACCAGTCTTCTTACATCGT | 7727-7753 |
| WHCV-R8 | TTTCACAAGTGCCGTGCCTAC | 9232-9252 |
| WHCV-F9 | GGTTTGCCTGGCACGATATTAC | 8883-8904 |
| WHCV-R9 | ACTTAGGTGTCTTAGGATTGGCTGTAT | 10345-10371 |
| WHCV-F10 | TTGTCATCTCGCAAAGGCTCT | 9974-9994 |
| WHCV-R10 | GAGATTATAAGAGCCCACATGGAAA | 11473-11497 |
| WHCV-F11 | GCTATGGGTATTATTGCTATGTCTGCT | 11124-11150 |
| WHCV-R11 | TGGATTTCCCACAATGCTGAT | 12557-12577 |
| WHCV-F12 | CTGATCAAGCTATGACCCAAATGT | 12295-12318 |
| WHCV-R12 | GCAACAGCTGGACAATCCTTAAGT | 13723-13746 |
| WHCV-F13 | TCTGCGGTATGTGGAAAGGTTAT | 13396-13418 |
| WHCV-R13 | GTCAGCAGCATACACAAGTAATTCCT | 14562-14587 |
| WHCV-F14 | AGGGCTTTAACTGCAGAGTCACAT | 14201-14224 |
| WHCV-R14 | GCGGACATACTTATCGGCAATT | 15598-15619 |
| WHCV-F15 | TCAATAGCCGCCACTAGAGGAG | 15188-15209 |
| WHCV-R15 | TCACCAGCATTTGTCCAGTCAC | 16587-16608 |
| WHCV-F16 | TTGGGGCTTGTGTTCTTTGC | 16257-16276 |
| WHCV-R16 | CAAGCAGGGTTACGTGTAAGGAAT | 17746-17769 |
| WHCV-F17 | TGTCAATGCCAGATTACGTGCT | 17410-17431 |
| WHCV-R17 | TAACAAAGCACTCGTGGACAGC | 18896-18917 |
| WHCV-F18 | TATGGGCACATGGCTTTGAGT | 18609-18629 |
| WHCV-R18 | TAAGAACACCATTACGGGCATTT | 20041-20063 |
| WHCV-F19 | TTGATGGACAACAGGGTGAAGTAC | 19680-19703 |
| WHCV-R19 | CGAAGTGTCCCATGAGCTTATAAA | 21213-21236 |

| | | |
|---|---|---|
| WHCV-F20 | AGGAGTTGCACCAGGTACAGCT | 20902-20923 |
| WHCV-R20 | ACCCACATAATAAGCTGCAGCAC | 22360-22382 |
| WHCV-F21 | CTATTAATTTAGTGCGTGATCTCCCTC | 22204-22230 |
| WHCV-R21 | AAATTTGTGGGTATGGCAATAGAGTTA | 23705-23731 |
| WHCV-F22 | ACTTACTCCTACTTGGCGTGTTTATTC | 23462-23488 |
| WHCV-R22 | GCATTAATGCCAGAGATGTCACC | 25077-25099 |
| WHCV-F23 | CTATCATCTTATGTCCTTCCCTCAGTC | 24716-24742 |
| WHCV-R23 | TAGTCGTCGTCGGTTCATCATAAAT | 26195-26219 |
| WHCV-F24 | TACTTCAGGTGATGGCACAACAA | 25915-25937 |
| WHCV-R24 | AAGCTCACAAGTAGCGAGTGTTATCA | 27435-27460 |
| WHCV-F25 | CGTGTAGCAGGTGACTCAGGTTT | 27094-27116 |
| WHCV-R25 | TACCGTCACCACCACGAATTC | 28567-28587 |
| WHCV-F26 | GGACCCCAAAATCAGCGAAAT | 28302-28322 |
| WHCV-R26 | AAAATCACATGGGGATAGCACTACT | 29840-29864 |

B. Primers for WHCV detection

| | | |
|---|---|---|
| S1423F | GCCGGTAGCACACCTTGTAA | 314bp |
| S1736R2 | GGATCACGGACAGCATCAGT | |
| S1869R1 | AGCAACAGGGACTTCTGTGC | |
| S2620F | ACTTCTGCACTGTTAGCGGG | 555bp |
| S3174R2 | ATGAGGTGCTGACTGAGGGA | |
| S3240R1 | GGCAGGAGCAGTTGTGAAGT | |

C. Primers for WHCV detection using qPCR
(designed based on the whole genome of WHCV (MN908947.3))

| | | |
|---|---|---|
| WHCV-F | TGATGATACTCTCTGACGATGCTGT | 15704-15728 |
| WHCV-R | CTCAGTCCAACATTTTGCTTCAGA | 15823-15846 |
| WHCV-P | ROX-ATGCATCTCAAGGTCTAGTG-MGB | 15749-15768 |

D. Primers used in 5'/3' RACE

| | | |
|---|---|---|
| 5-GSP | CCACATGAGGGACAAGGACACCAAGTG | 573-599 (599bp) |
| 5-GSPn | CATGACCATGAGGTGCAGTTCGAGC | 491-515 (515bp) |
| 3-GSP | TGTCGCGCATTGGCATGGAAGTCACACC | 29212-29239 (688bp) |
| 3-GSPn | CTCAAGCCTTACCGCAGAGACAGAAG | 29398-29423 (502bp) |

E. Primers for detection of other respiratory pathogens using qPCR

| | |
|---|---|
| 1012FluA-Fv1 | GGARTGGMTAAAGACAAGACCAATC |
| 1012FluA-Rv1 | GGCRTTYTGGACAAASCGTCTAC |
| 1012FluA-Pv1 | ROX-AGTCCTCGCTCACTGGGCACGGT-BHQ2 |
| 1083FluB-Fv* | AGACCAGAGGGAAACTATGCCC |
| 1083FluB-Rv* | TCCGGATGTAACAGGTCTGACTT |
| 1083FluB-Pv*(Victoria) | CY5-CAGACCAAAATGCACGGGGAAHATACC-BHQ1 |
| 1083FluB-Pv*(Yamagata) | FAM-CAGRCCAATGTGTGTGGGGAYCACACC-BHQ1 |

| | |
|---|---|
| 1111HADV-Fv1 | GCCACGGTGGGGTTTCTAAACTT |
| 1111HADV-Rv1 | GCCCCAGTGGTCTTACATGCACATC |
| 1111HADV-Pv1 | FAM-TGCACCAGACCCGGGCTCAGGTACTCCGA-TAMRA |
| 1281CPn-Fv3 | AGCACAAACACCTCAGACTACAC |
| 1281CPn-Rv3 | AGAACAATGCCGATTCCTAAG |
| 1281CPn-Pv3 | FAM-ACAACCATCAGTATCTCACAAGGCAACAC-BHQ1 |

### Supplementary Figures

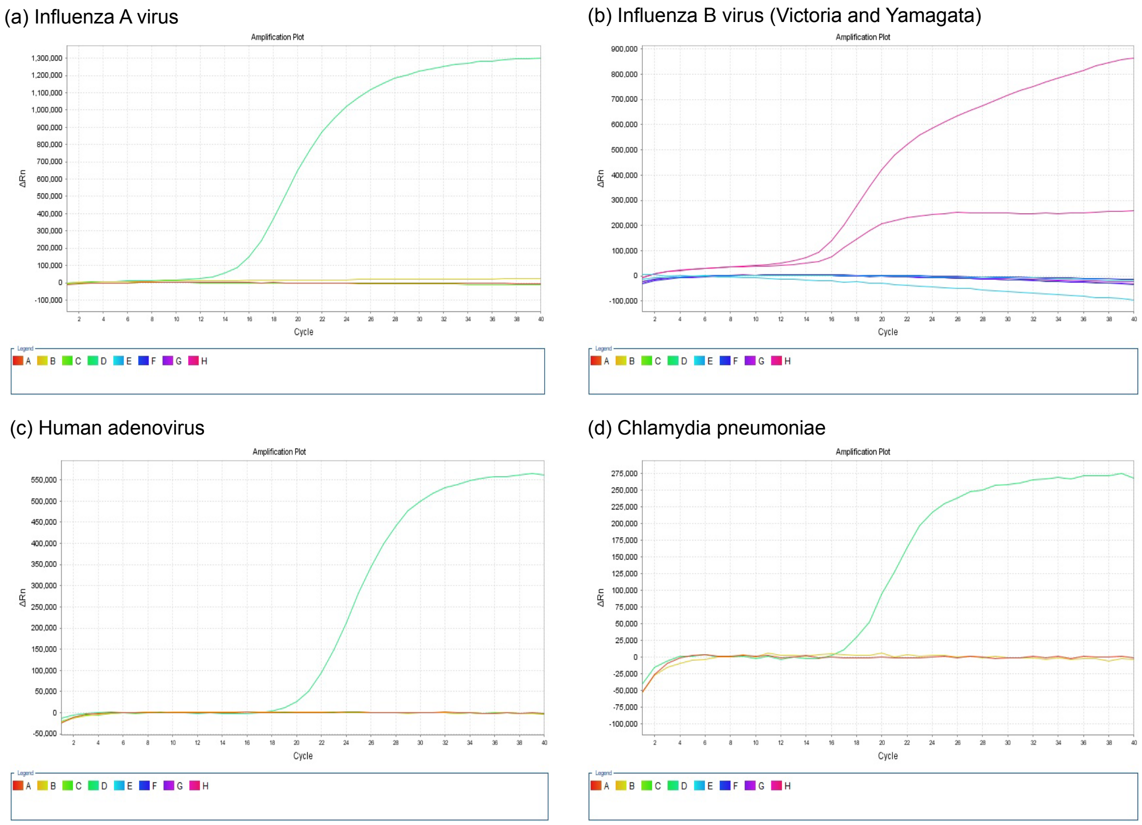

**Figure S1.** Detection of other respiratory pathogens by qPCR.

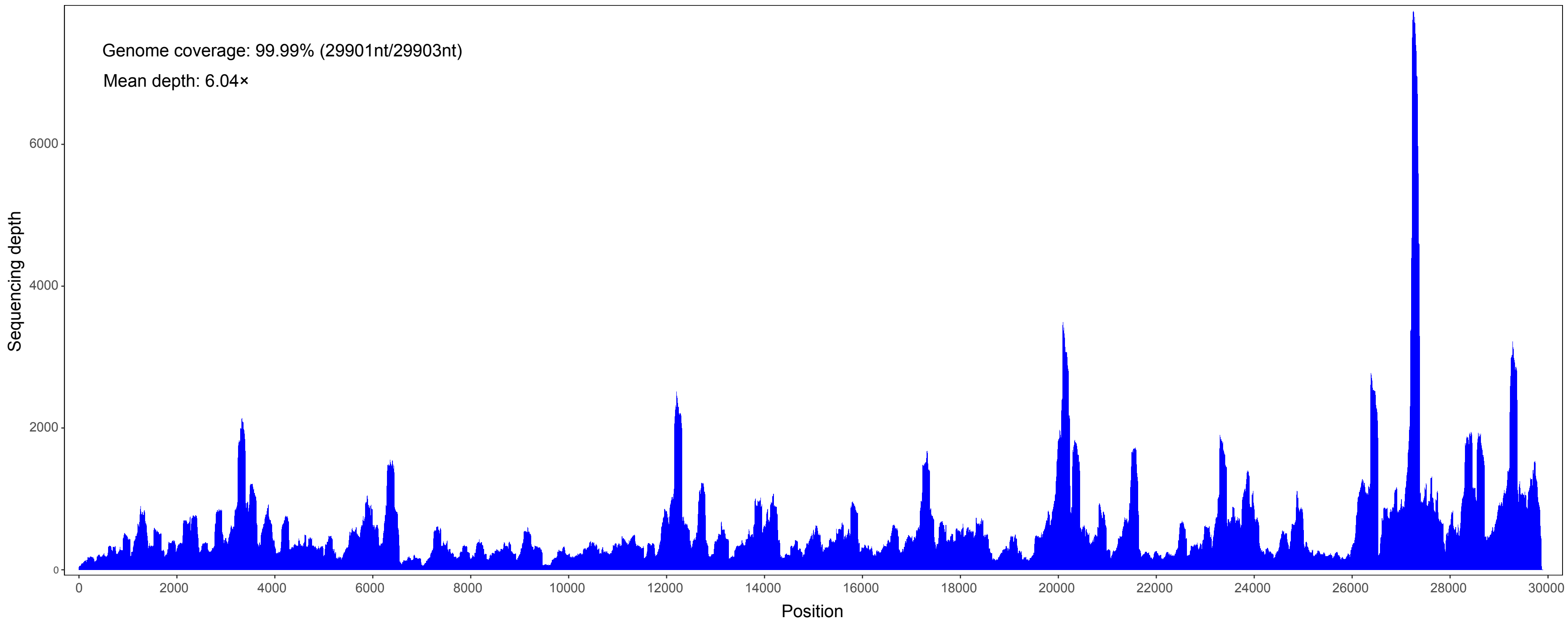

**Figure S2.** Mapped read count plot showing the coverage depth per base of the WHCV genome.

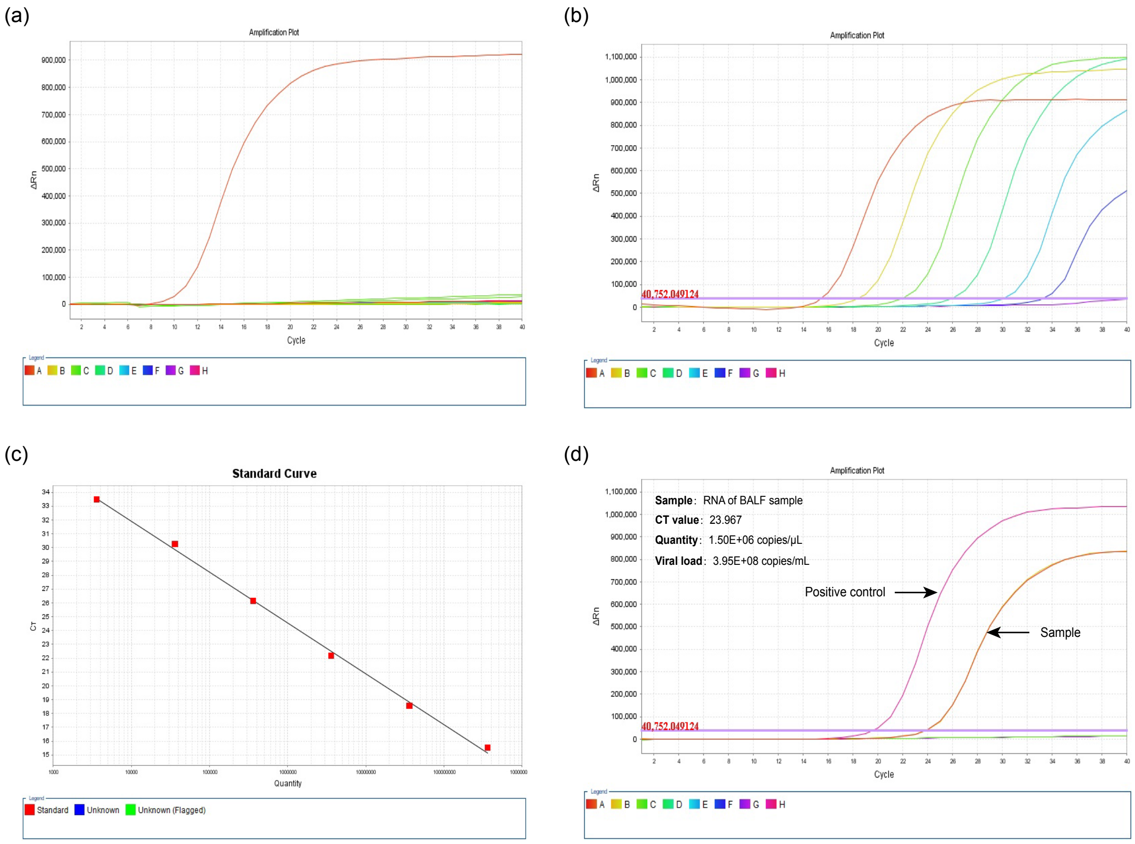

**Figure S3.** Detection of WHCV in clinical samples by RT-qPCR. **(a)** Specificity of the WHCV primers used in RT-qPCR. Test samples comprised clinical samples that are positive for at least one of the following viruses: Influenza A virus (09H1N1 and H3N2), Influenza B virus, Human adenovirus, Respiratory syncytial virus, Rhinovirus, Parainfluenza virus type 1-4, Human bocavirus, Human metapneumovirus, Coronavirus OC43, Coronavirus NL63, Coronavirus 229E and Coronavirus HKU1. **(b-c)** Standard curve. **(d)** Amplification curve of WHCV.

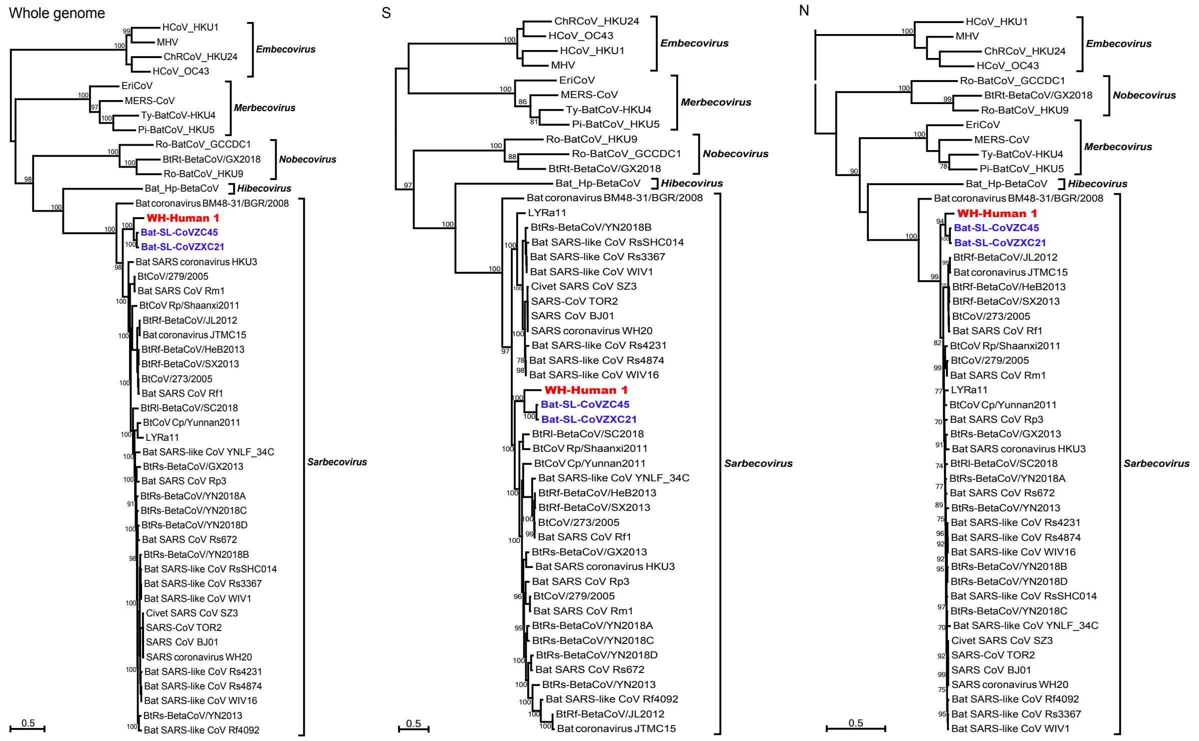

**Figure S4**. Maximum likelihood phylogenetic trees of the nucleotide sequences of the whole genome, S and N genes of WHCV and related coronaviruses. Numbers (>70) above or below branches indicate percentage bootstrap values. The trees were mid-point rooted for clarity only. The scale bar represents the number of substitutions per site.

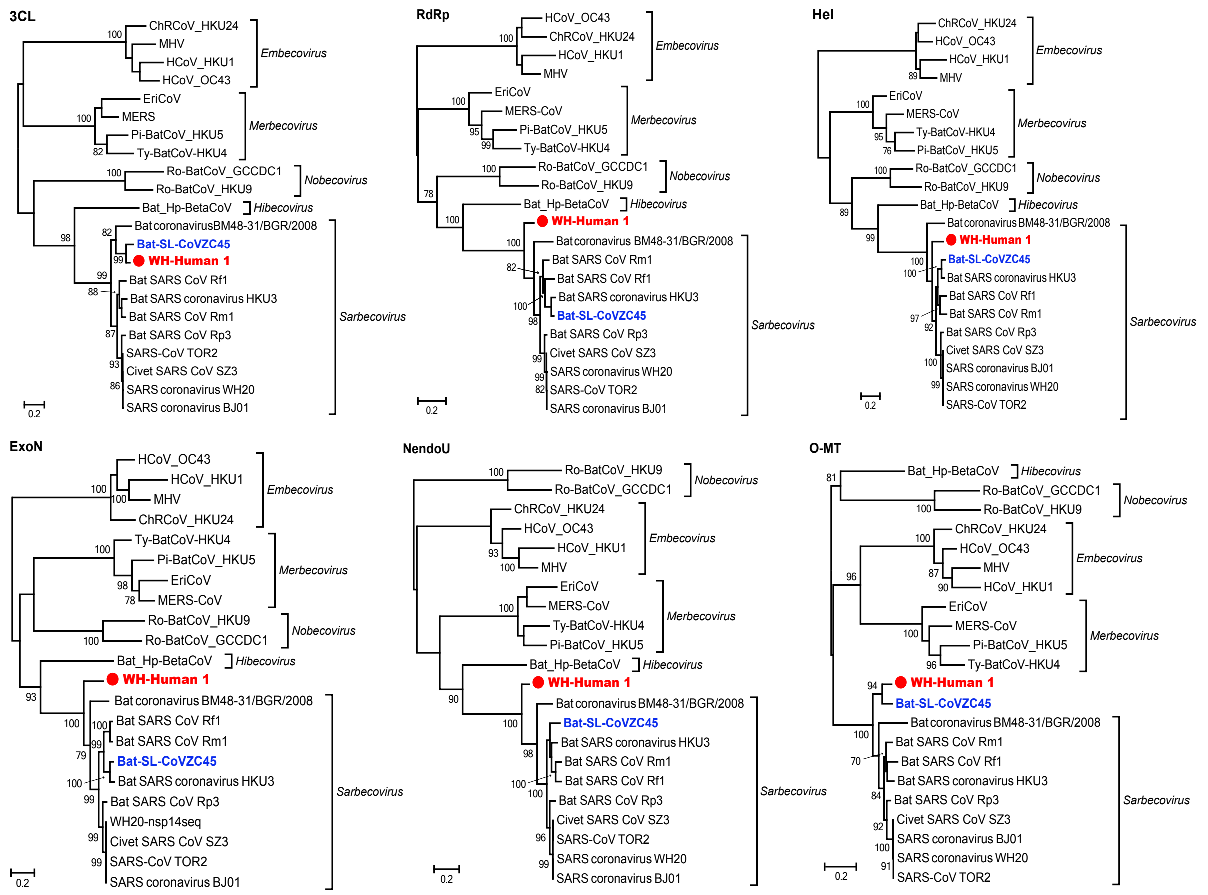

**Figure S5**. Maximum likelihood phylogenetic trees of the nucleotide sequences of the 3CL, RdRp, Hel, ExoN, NendoU, and O-MT genes of WHCV and related coronaviruses. Numbers (>70) above or below branches indicate percentage bootstrap values. The trees were mid-point rooted for clarity only. The scale bar represents the number of substitutions per site.

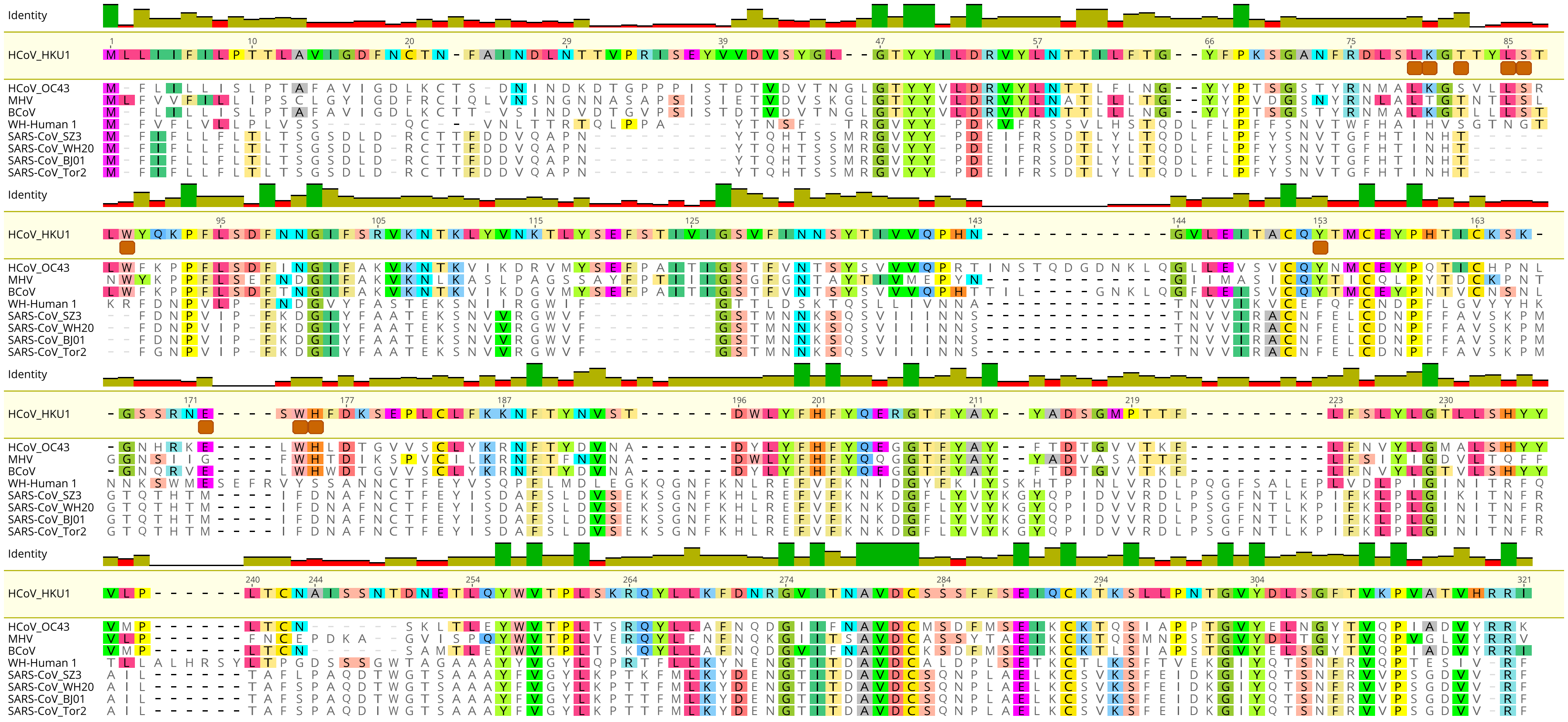

**Figure S6.** Amino acid sequence comparison of the N-terminal domain (NTD) of spike protein of WHCV, bovine coronavirus (BCoV), mouse hepatitis virus (MHV) and human coronavirus (HCoV OC43 and HKU1) that can bind to sialic acid and the SARS-CoVs that cannot. The key residues[13] for sialic acid binding on BCoV, MHV, HCoV OC43 and HKU1 were marked with a brown box.

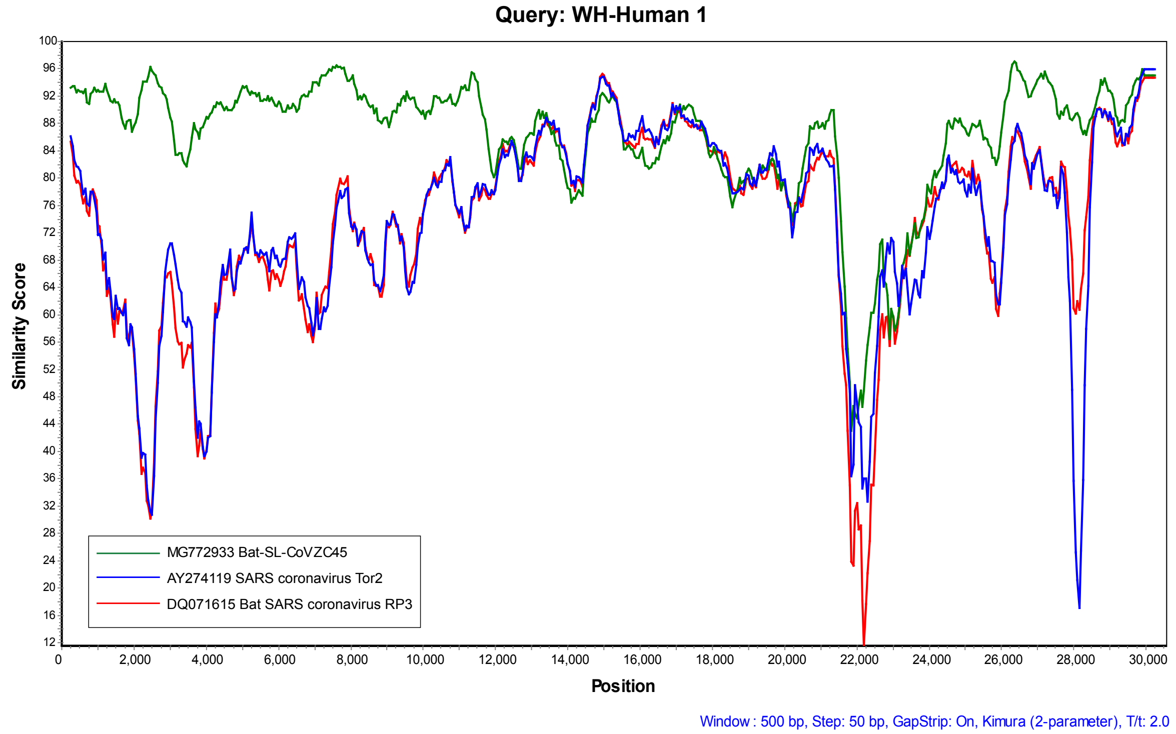

**Figure S7.** A sequence similarity plot of WHCV, SARS- and bat SARS-like CoVs revealing putative recombination events.
